# Larger but younger fish when growth outpaces mortality in heated ecosystem

**DOI:** 10.1101/2022.04.13.488128

**Authors:** Max Lindmark, Malin Karlsson, Anna Gårdmark

## Abstract

Ectotherms are predicted to “shrink” with global warming, in line with general growth models and the temperature-size rule (TSR), both predicting smaller adult sizes with warming. However, they also predict faster juvenile growth rates and thus larger size-at-age of young organisms. Hence, the result of warming on the size-structure of a population depends on the interplay between how mortality rate, juvenile- and adult growth rates are affected by warming. Here, we use two-decade long time series of biological samples from a unique enclosed bay heated by cooling water from a nearby nuclear power plant to become 5–10 °C warmer than its reference area. We used growth-increment biochronologies (12658 reconstructed length-at-age estimates from 2426 individuals) to quantify how >20 years of warming has affected body growth and size-at-age and catch data to quantify mortality rates and population size- and age structure of Eurasian perch (*Perca fluviatilis*). In the heated area, growth rates were faster for all sizes, and hence size-at-age was larger for all ages, compared to the reference area. While mortality rates were also higher (lowering mean age by 0.4 years), the faster growth rates lead to a 2 cm larger mean size in the heated area. Differences in the size-spectrum exponent (describing the proportion of fish by size) were less clear statistically. Our analyses reveal that mortality, in addition to plastic growth and size-responses, is a key factor determining the size structure of populations exposed to warming. Understanding the mechanisms by which warming affects the size- and age structure of populations is critical for predicting the impacts of climate change on ecological functions, interactions, and dynamics.

## Introduction

Ectotherm species, constituting 99% of species globally (Atkinson and Sibly, 1997; Wilson, 1992), are commonly predicted to shrink in a warming world (Gardner et al., 2011; Sheridan and Bickford, 2011). However, as the size-distribution of many species spans several orders of magnitude, and temperature-effects on size may vary with size or age, it is important to be specific about which sizes- or life stages are predicted to shrink (usually mean or adult is meant). For instance, warming can shift size-distributions without altering mean size if increases in juvenile size-at-age outweigh the decline in size-at-age in adults, which is consistent with the temperature size-rule, TSR (Atkinson, 1994). Resolving how warming induces changes in population size-distributions may thus be more instructive (Fritschie and Olden, 2016), especially for inferring warming effects on species’ ecological role, biomass production, or energy fluxes (Gårdmark and Huss, 2020; Yvon-Durocher et al., 2011). This is because key processes such as metabolism, feeding, growth, mortality scale with body size (Andersen, 2020; Blanchard et al., 2017; Brown et al., 2004; Pauly, 1980; Thorson et al., 2017; Ursin, 1967). Hence, as the value of these traits at mean body size is not the same as the mean population trait value (Bernhardt et al., 2018), the size-distribution within a population matters for its dynamics and for how it changes under warming.

The population size distribution can be represented as a size-spectrum, which generally is the frequency distribution of individual body sizes (Edwards et al., 2017). It is often described in terms of the size-spectrum slope (slope of individuals or biomass of a size class over the mean size of that class on log-log scale (Edwards et al., 2017; Sheldon et al., 1973; White et al., 2007)) or simply the exponent of the power law individual size-distribution (Edwards et al., 2017). The size-spectrum thus results from temperature-dependent ecological processes such as body growth, mortality and recruitment (Blanchard et al., 2017; Heneghan et al., 2019). Despite its rich theoretical foundation (Andersen, 2019) and usefulness as an ecological indicator (Blanchard et al., 2005), few studies have evaluated warming-effects on the species size-spectrum in larger bodied species (but see Blanchard et al., 2005), and none in large scale experimental set-ups. There are numerous paths by which a species’ size-spectrum could change with warming (Heneghan et al., 2019). For instance, in line with TSR predictions, warming may lead to a smaller size-spectrum exponents (steeper slope) if the maximum size declines. However, changes in size-at-age and the relative abundances of juveniles and adults may alter this decline in the size-spectrum slope. Warming can also lead to elevated mortality (Barnett et al., 2020; Berggren et al., 2021; Biro et al., 2007; Pauly, 1980), partly because a faster pace of life with higher metabolic rates are associated with shorter lifespan (Brown et al., 2004; Munch and Salinas, 2009) or due to direct lethal effects of extreme temperature events. This truncates the age-distribution towards younger individuals (Barnett et al., 2017). This may reduce density dependence and potentially increase growth rates, thus countering the effects of mortality on the size spectrum exponent. However, not all sizes may benefit from warming, as e.g. the optimum temperature for growth declines with size (Lindmark et al., 2022). Hence, the effect of warming on the size-spectrum depends on several interlinked processes affecting abundance-at-size and size-at-age.

Size-at-age is generally predicted to increase with warming for small individuals, but decrease for large individuals according to the mentioned TSR (Atkinson, 1994; Ohlberger, 2013). Several factors likely contribute to this pattern, such as increased allocation to reproduction (Wootton et al., 2022) and larger individuals in fish populations having optimum growth rates at lower temperatures (Lindmark et al., 2022). Empirical support in fishes for this pattern seem to be more consistent for increases in size-at-age of juveniles (Huss et al., 2019; Rindorf et al., 2008; Thresher et al., 2007) than declines in adult size-at-age (but see (Baudron et al., 2014; Oke et al., 2022; Smoliński et al., 2020)), for which a larger diversity in responses is observed among species (Barneche et al., 2019; e.g., Huss et al., 2019). However, most studies have been done on commercially exploited species, since long time series are more common in such species. This may confound or interact with effects of temperature, because fishing mortality can affect density dependent growth (van Gemert and Andersen, 2018), but also select for slow growing individuals and changes in maturation processes, which also influences growth trajectories (Audzijonyte et al., 2016).

The effect of temperature on mortality rates of wild populations are more often studied using among-species analyses (Pauly, 1980; Thorson et al., 2017). These relationships based on thermal gradients in space may not necessarily be the same as the effects of *warming* on mortality on single populations. Hence, the effects of warming on growth and size-at-age and mortality within natural populations constitute a key knowledge gap for predicting the consequences of climate change on population size-spectra.

Here we used data from a unique, large-scale 23-year-long heating-experiment of a coastal ecosystem to quantify how warming changed fish body growth, mortality, and the size structure in an unexploited population of Eurasian perch (*Perca fluviatilis*, ‘perch’). We compare fish from this enclosed bay exposed to temperatures approximately 5–10 °C above normal (‘heated area’) with fish from a reference area in the adjacent archipelago (Fig. 1). Using hierarchical Bayesian models, we quantify differences in key individual- and population level parameters, such as body growth, asymptotic size, mortality rates, and size-spectra, between the heated and reference coastal area.

**Fig. 1.**
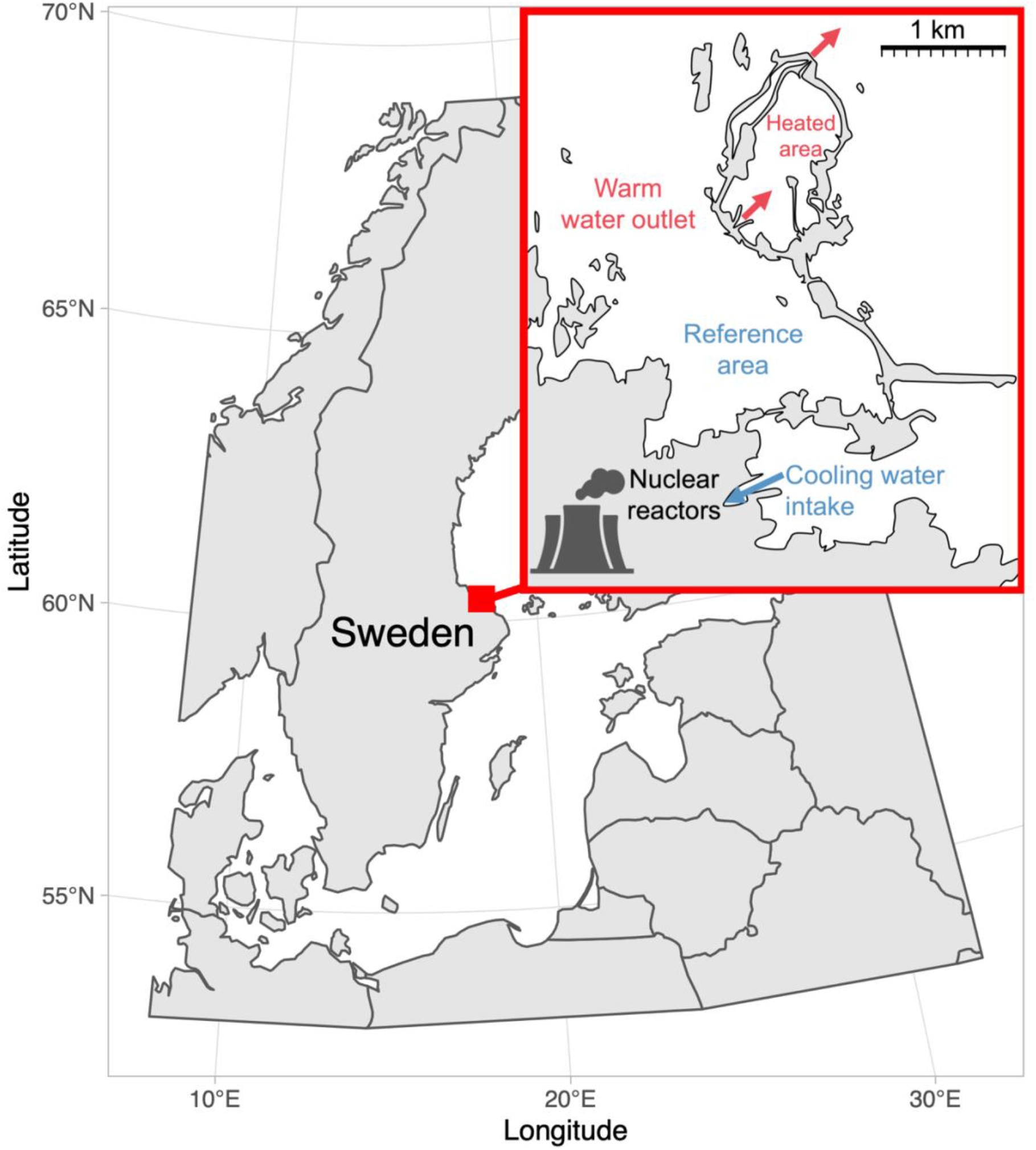
Map of the area with the unique whole-ecosystem warming experiment from which perch in this study was sampled. Inset shows the 1 km^2^ enclosed coastal bay that has been artificially heated for 23 years, the adjacent reference area with natural temperatures, and locations of the cooling water intake and where the heated water outlet from nuclear power plants enters the heated coastal basin. The arrows indicate the direction of water flow.

## Materials and Methods

### Data

We use size-at-age data from perch sampled annually from an artificially heated enclosed bay (‘the Biotest basin’) and its reference area, both in the western Baltic Sea (Fig. 1). Heating started in 1980, the first analyzed cohort is 1981, and first and last catch year is 1987 and 2003, respectively, to omit transient dynamics and acute responses, and to ensure we use cohorts that only experienced one of the thermal environments during its life. A grid at the outlet of the heated area (Fig. 1) prevented fish larger than 10 cm from migrating between the areas (Adill et al., 2013; Huss et al., 2019), and genetic studies confirm the reproductive isolation between the two populations during this time-period (Björklund et al., 2015). However, the grid was removed in 2004, and since then fish growing up in the heated Biotest basin can easily swim out, fish caught in the reference area cannot be assumed to be born there. Hence, we use data only up until 2003. This resulted in 12658 length-at-age measurements from 2426 individuals (i.e., multiple measurements per individual) from 256 net deployments.

We use data from fishing events using survey-gillnets that took place in October in the heated Biotest basin and in August in the reference area when temperatures are most comparable between the two areas (Huss et al., 2019), because temperature affects catchability in static gears. The catch was recorded by 2.5 cm length classes during 1987-2000, and into 1 cm length groups 2001-2003. To express lengths in a common length standard, 1 cm intervals were converted into 2.5 cm intervals. The unit of catch data is hence the number of fish caught by 2.5 cm size class per net per night (i.e., a catch-per-unit-effort [CPUE] variable). All data from fishing events with disturbance affecting the catch (e.g., seal damage, strong algal growth on the gears, clogging by drifting algae) were removed (years 1996 and 1999 from the heated area in the catch data).

Length-at-age throughout an individuals’ life was reconstructed for a random or length-stratified subset of caught individuals each year (depending on which year, and in some cases, the number of fish caught). This was done using growth-increment biochronologies derived from annuli rings on the operculum bones (with control counts done on otoliths). Such analyses have become increasingly used to analyze changes in growth and size-at-age of fishes (Essington et al., 2022; Morrongiello and Thresher, 2015). Specifically, an established power-law relationship between the distance of annual rings and fish length was used: *L* = *κR^s^*, where *L* is the length of the fish, *R* the operculum radius, *κ* the intercept, and s the slope of the line for the regression of log-fish length on log-operculum radius from a large reference data set for perch (Thoresson, 1996). Back-calculated length-at-age were obtained from the relationship 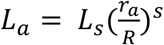, where *L_a_* is the back-calculated body length at age *a, L_s_* is the final body length (body length at catch), *r_a_* is the distance from the center to the annual ring corresponding to age *a* and *s* = 0.861 for perch (Thoresson, 1996). Since perch exhibits sexual size-dimorphism, and age-determination together with back calculation of growth was not done for males in all years, we only used females for our analyses.

### Statistical Analysis

The differences in size-at-age, growth, mortality, and size structure between perch in the heated and the reference area were quantified using hierarchical linear and non-linear models fitted in a Bayesian framework. First, we describe each statistical model and then provide details of model fitting, model diagnostics and comparison.

To describe individual growth throughout life, we fit the von Bertalanffy growth equation (VBGE) (Beverton and Holt, 1957; von Bertalanffy, 1938) on a log scale, describing length as a function of age to evaluate differences in size-at-age and asymptotic size: log(*L_t_*) = log(*L*_∞_(1 – *e*^(-*K*(*t*–*t*_0_))^)), where *L_t_* is the length-at-age (*t*, years), *L*_∞_ is the asymptotic size, *K* is the Brody growth coefficient (*yr*^−1^) and *t*_0_ is the age when the average length was zero. Here and henceforth, log refers to natural logarithms. We used only age- and size-at-catch as the response variables (i.e., not back-calculated length-at-age). This was to have a simpler model and not have to account for parameters varying within individuals as well as cohorts, as mean sample size per individual was only ~5. We let parameters vary among cohorts rather than year of catch, because individuals within cohorts share similar environmental conditions and density dependence (Morrongiello and Thresher, 2015). Eight models in total were fitted (with area being dummy-coded), with different combinations of shared and area-specific parameters. We evaluated if models with area-specific parameters led to better fit and quantified the differences in area-specific parameters (indexed by subscripts heat and ref). The model with all area-specific parameter can be written as:

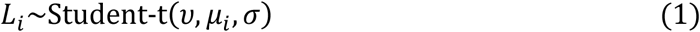

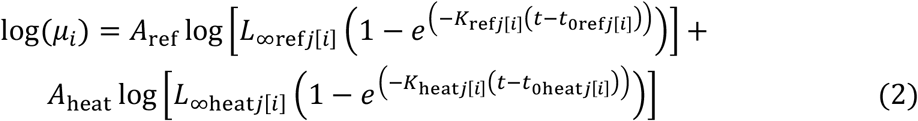

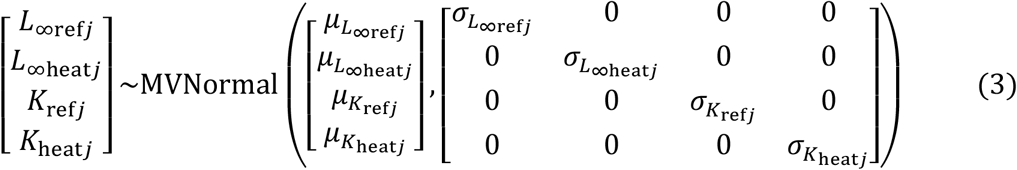

where lengths are Student-t distributed to account for extreme observations, *ν, μ* and *ϕ* represent the degrees of freedom, mean and the scale parameter, respectively. *A*_ref_ and *A*_heat_ are dummy variables such that *A*_ref_ = 1 and *A*_heat_ = 0 if it is the reference area, and vice versa for the heated area. The multivariate normal distribution in Eq. 3 is the prior for the cohort-varying parameters *L*_∞ref*j*_, *L*_∞heat*j*_, *K*_ref*j*_ and *K*_heat*j*_ (for cohorts *j* = 1981,…,1997) (note that cohorts extend further back in time than the catch data), with hyper-parameters *μ*_*L*_∞ref__, *μ*_*L*_∞heat__, *μ*_*K*_ref__, *μ*_*K*_heat__ describing the non-varying population means and a covariance matrix with the between-cohort variation along the diagonal (note we did not model a correlation between the parameters, hence off-diagonals are 0). The other seven models include some or all parameters as parameters common for the two areas, e.g., substituting *L*_∞ref*j*_ and *L*_∞heat*j*_ with *L*_∞*j*_. To aid convergence of this non-linear model, we used informative priors chosen after visualizing draws from prior predictive distributions (Wesner and Pomeranz, 2021) using probable parameter values (*Supporting Information*, Fig. S1, S11). We used the same prior distribution for each parameter class for both areas to not introduce any other sources of differences in parameter estimates between areas. We used the following priors for the VBGE model: *μ*_L_∞ref,heat__~N(45,20), *μ*_*K*_ref,heat__~N(0.2, 0.1), *t*_0ref,heat_~N(−0.5,1) and *ν*~gamma(2,0.1). *σ* parameters, *μ*_*L*_∞ref__, *μ*_*L*_∞heat__, *μ*_K_ref__, *μ*_K_heat__ were given a Student-t(3,0,2.5) prior.

We also compared how body growth scales with body size (in contrast to length vs age). This is because size-at-age reflects lifetime growth history rather than current growth histories and may thus be large because growth was fast early in life, not because current growth rates are fast (Lorenzen, 2016). We therefore fit allometric growth models describing how specific growth rate scales with length: *G* = *αL^θ^*, where *G*, the annual specific growth between year *t* and *t* + 1, is defined as: *G* = 100 × (log (*L*_*t*+1_) – *log* (*L_t_*)) and *L* is the geometric mean length: *L* = (*L*_*t*+1_ × *L_t_*)^0.5^. Here we also used back-calculated length-at-age, resulting in multiple observations for each individual. As with the VBGE model, we dummy coded area to compare models with different combinations of common and shared parameters. We assumed growth rates were Student-t distributed, and the full model can be written as:

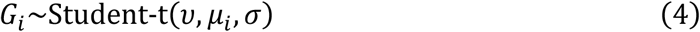

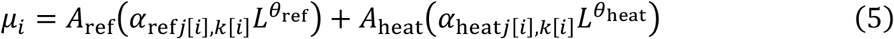

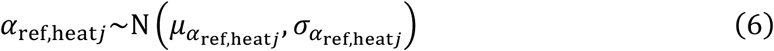

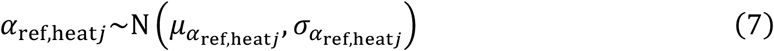

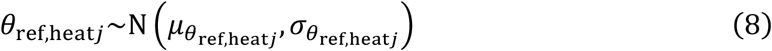

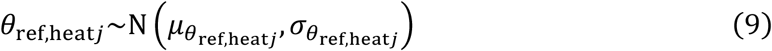

We assumed only *α* varied across individuals *j* within cohorts *k* and compared two models: one with *θ* common for the heated and reference area, and one with an area-specific *θ*. We used the following priors: *α*_ref,heat_~N(500,100), *θ*_ref,heat_~N(−1.2,0.3) and *ν*~gamma(2,0.1). *σ*, *σ*_id;cohort_ and *σ*_cohort_ were all given a Student-t(3,0,13.3) prior.

We estimated total mortality by fitting linear models to the natural log of catch (CPUE) as a function of age (catch curve regression), under the assumption that in a closed population, the exponential decline can be described as *N_t_* = N_0_*e*^−*Zt*^, where *N_t_* is the population at time *t*, *N*_0_ is the initial population size and *Z* is the instantaneous mortality rate. This equation can be rewritten as a linear equation: log(*C_t_*) = log(*νN*_0_) – *Zt*, where *C_t_* is catch at age *t*, if catch is assumed proportional to the number of fish (i.e., *C_t_* = *νN_t_*. Hence, the negative of the slope of the regression is the mortality rate, *Z*. To get catch-at-age data, we constructed area-specific age-length keys using the sub-sample of the total (female) catch that was age-determined. Age length-keys describe the age-proportions of each length-category (i.e., a matrix with length category as rows, ages as columns). Age composition is then estimated for the total catch based on the “probability” of fish in each length-category being a certain age. Lastly, due to the smallest and youngest fish not being representatively caught with the gillnet, catch is dome shaped over size and age. We therefore followed the practice of selecting only ages on the descending right limb (Dunn et al., 2002) (*Supporting Information*, Fig. S14). With fit this model with and without an *age* × *area*-interaction, and the former can be written as:

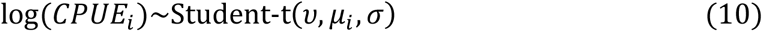

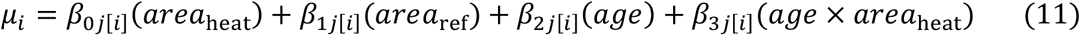

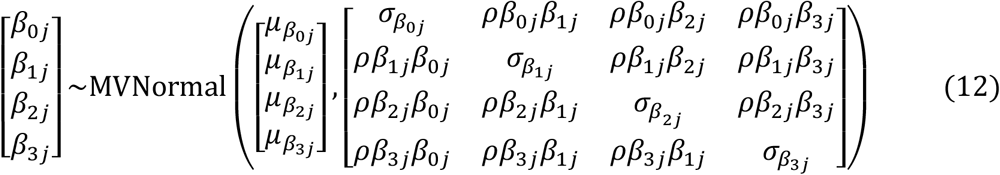

where *β*_0*j*_ and *β*_1*j*_ are the intercepts for the reference and heated areas, respectively, *β*_2*j*_ is the age slope for the reference area and *β*_3*j*_ is the difference between the age slope in the reference area and in the heated area. All parameters vary by cohort (for cohort *j* = 1981,…, 2000). We use the default *brms* priors for this models, i.e., flat priors for the regression coefficients (Bürkner, 2017) and *ν*~gamma(2,0.1). *σ* and σ_β_0,…,3__ were given a Student-t(3,0,2.5) prior.

Lastly, we quantified differences in the average age and size-distributions between the areas. We estimate the biomass size-spectrum exponent *γ* directly, using the likelihood approach for binned data, i.e., the *MLEbin* method in the R package *sizeSpectra* (Edwards, 2020; Edwards et al., 2020, 2017). This method explicitly accounts for uncertainty in body masses *within* size-classes (bins) in the data and has been shown to be less biased than regression-based methods or the likelihood method based on bin-midpoints (Edwards et al., 2020, 2017). We pooled all years to ensure negative relationships between biomass and size in the size-classes (as the sign of the relationship varied between years). We also fitted lognormal models (due to strictly positive values presence of a tail) to length- and age resolved catch data. Here we assume that the catchability with respect to size does not differ between the areas, and therefore use the entire catch as data (*Supporting Information*, Fig. S16) (note, in contrast to the catch curve regression, we do not need to filter representatively caught size or age classes, because we do include an age or size). The lognormal models fitted to age or size (denoted *y_age,length,i_*) model can be written as:

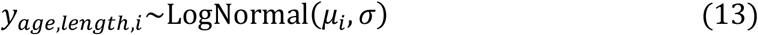

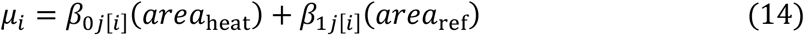

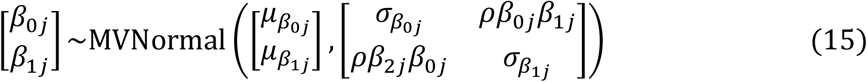

where *β*_0*j*_ is the intercept for the reference area and *β*_1*j*_ is the intercept for the heated area. These intercepts vary by year (for years *j* = 1987,…, 2003). We use flat priors for the regression coefficients, and *σ* was given a Student-t(3,0,2.5) prior and compared models with and without random slopes.

All analyses were done using R (R Core Team, 2020) version 4.0.2 with R Studio (2021.09.1). The packages within the *tidyverse* (Wickham et al., 2019) collection were used to processes and visualize data. Models where fit using the R package *brms* (Bürkner, 2018). For the non-linear von Bertalanffy growth equation and the allometric growth model, we used informative priors to facilitate convergence. These were chosen by defining vague priors, and the progressively tightening these until convergence was achieved (Bürkner, 2017; Gesmann and Morris, 2020). We used prior predictive checks to ensure the priors were suitable (vague enough to include also very unlikely prediction, but informative enough to ensure convergence), and the final prior predictive checks are shown in Figs. S1 and S11. We also explored priors vs posteriors to evaluate the influence of our informative priors visually (*Supporting Information*, Figs. S7 and S13). For the linear models (catch curve and mean size), which do not require the same procedure to achieve convergence typically, we used the default priors from *brms* as written above. We used 3 chains and 4000 iterations in total per chain. Models were compared by evaluating their expected predictive accuracy (expected log pointwise predictive density) using leave-one-out cross-validation (LOO-CV) (Vehtari et al., 2017) while ensuring Pareto *k* values < 0.7, in the R package *loo* (Vehtari et al., 2020). Results of the model comparison can be found in the *Supporting Information*, Table S1-S2. We used *bayesplot* (Gabry et al., 2019) and *tidybayes* (Kay, 2019) to process and visualize model diagnostics and posteriors. Model convergence and fit was assessed by ensuring potential scale reduction factors 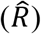 were less than 1.1, suggesting all three chains converged to a common distribution) (Gelman et al., 2003), and by visually inspecting trace plots, residuals QQ-plots and with posterior predictive checks (*Supporting Information*, Fig. S2, S12, S15, S17).

## Results

Analysis of fish (perch) size-at-age using the von Bertalanffy growth equation (VBGE) revealed that fish cohorts (year classes) in the heated area both grew faster initially (larger size-at-age) and reached larger predicted asymptotic sizes than those in the unheated reference area (Fig. 2). The model with area-specific VBGE parameters (*L*_∞_, *K* and *t*_0_) had best out of sample predictive accuracy (the largest expected log pointwise predictive density for a new observation; Table S1), and there is a clear difference in the predicted length-at-age (Fig. 2). Models where both *L*_∞_ and *K* were shared did not converge (*Supporting Information*, Table S1). Both the estimated values for fish asymptotic length (*L*_∞_) and growth coefficient (*K*) were larger in the heated compared to the reference area (*Supporting Information*, Fig. S8). We estimated that the asymptotic length of fish in the heated area was 16% larger than in the reference area (calculated as 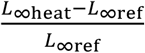) (*L*_∞heat_ = 45.7[36.8,56.3], L_∞ref_ = 39.4[35.4,43.9], where the point estimate is the posterior median and values in brackets correspond to the 95% credible interval). The growth coefficient was 27% larger in the heated area (*K*_heat_ = 0.19[0.15,0.23], *K*_ref_ = 0.15[0.12,0.17]). These differences in growth parameters lead to fish being approximately 11% to 7% larger in the heated area at any age relative to the reference area (*Supporting Information*, Fig. S4). Due to the last three cohorts (1995-1997) having large estimates of L∞_heat_ and low *K* (potentially due to their negative correlation and because of the young average age with data far from the asymptote, *Supporting Information*, Fig. S3 and S5-S6), we fit the same model with these cohorts omitted to evaluate the importance of those for the predicted difference between the areas. Without these, the predicted difference in size-at-age was still clear, but smaller (between 7% to 4%, *Supporting Information*, Fig. S9-10).

**Fig. 2.**
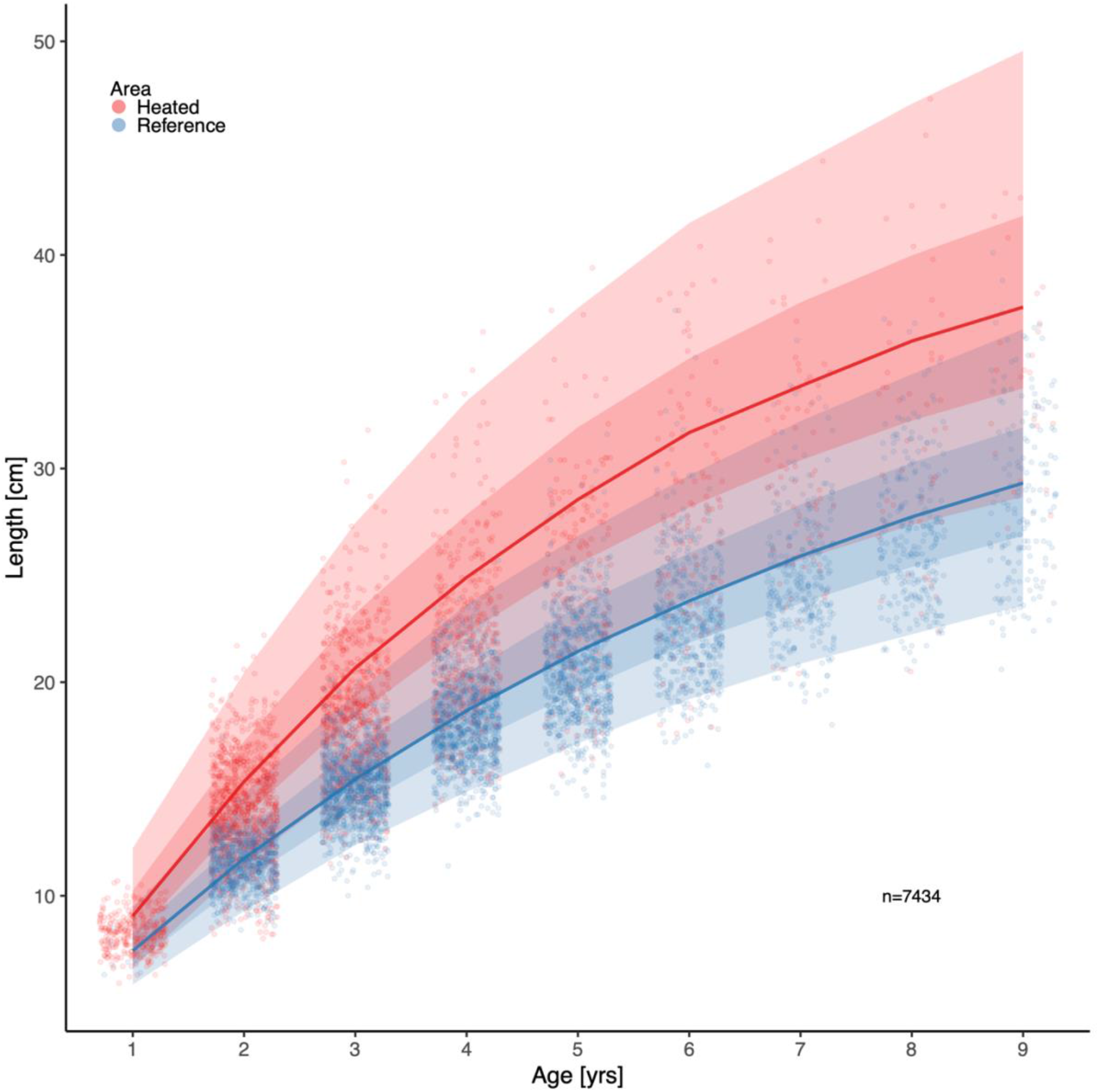
Fish grow faster and reach larger sizes in the heated (red) enclosed bay compared to the reference (blue) area. Points depict individual-level length-at-age and lines show the posterior draws of the global posterior predictive distribution (without group-level effects), both exponentiated, from the von Bertalanffy growth model with area-specific coefficients. The shaded areas correspond to 50% and 90% credible intervals.

In addition, we found that growth rates in the reference area were both slower and declined faster with size compared to the heated area (Fig. 3). The best model for growth (*G* = *αL^θ^*) had area-specific *α* and *θ* parameters (Table S2). Initial growth (*α*) was estimated to be 18% faster in the heated than in the reference area (*α*_heat_ = 512[462,565], *α*_ref_ = 433[413,454]), and growth of fish in the heated area declines more slowly with length than in the reference area (*θ*_heat_ = −1.13[−1.16, −1.11], *θ*_ref_ = −1.18[−1.19, −1.16]). The distribution of differences of the posterior samples for *α* and *θ* only had 0.3% and 0.2%, respectively, of the density below 0 (Fig. 3C, E), indicating high probability that length-based growth rates are faster in the heated area.

**Fig. 3.**
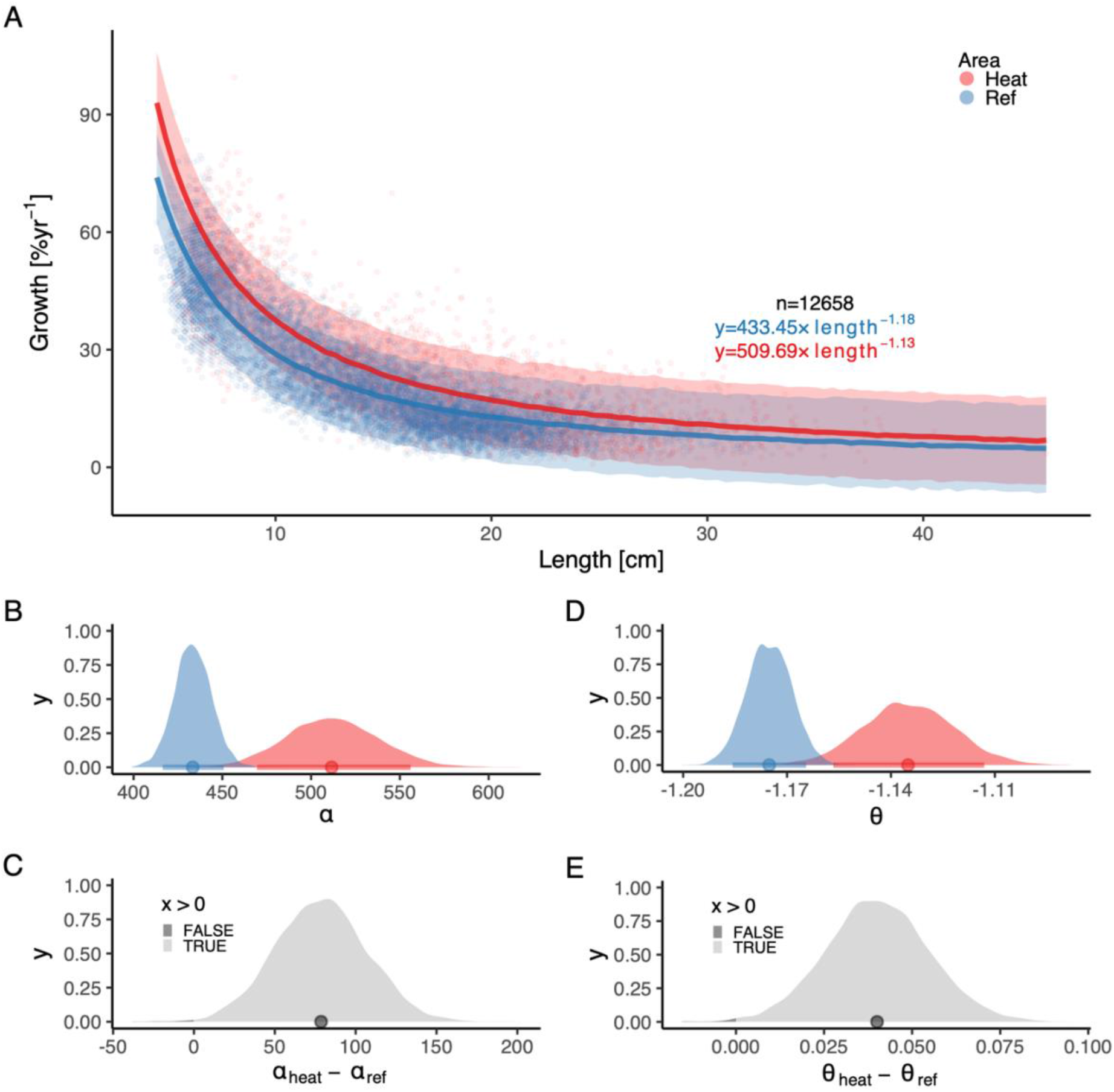
The faster growth rates in the heated area (red) compared to the reference (blue) are maintained as fish grow. The points illustrate specific growth estimated from back-calculated length-at-age (within individuals) as a function of length (expressed as the geometric mean of the length at the start and end of the time interval). Lines show the posterior draws of the global posterior predictive distribution (without group-level effects) from the allometric growth model with area-specific coefficients. The shaded areas correspond to the 90% credible interval. The equation uses mean parameter estimates. Panel (B) shows the posterior distributions for initial growth (*α*_heat_ (red) and *α*_ref_ (blue)), and (C) the distribution of their difference. Panel (D) shows the posterior distributions for the allometric decline in growth with length (*θ*_heat_ and *θ*_ref_), and (E) the distribution of their difference. The fill color depicts the area below 0 (0.3% and 0.2% for *α* and *θ*, respectively).

By analyzing the decline in catch-per-unit-effort over age, we found that the instantaneous mortality rate *Z* (rate at which log abundance declines with age) is higher in the heated area (Fig. 4). *Z* was estimated as a fixed effect, as the model where only intercepts varied among years had the best out of sample predictive ability. The overlap with zero is 0.07% for the distribution of differences of posterior samples of *Z*_heat_ and *Z*_ref_ (Fig. 4C). We estimated *Z*_heat_ to be 0.73[0.66,0.79] and *Z*_ref_ to be 0.62[0.58,0.67], which corresponds to annual mortality rates (calculated as 1 – *e*^−*z*^) of 52% in the heated area and 47% in the reference area.

**Fig. 4.**
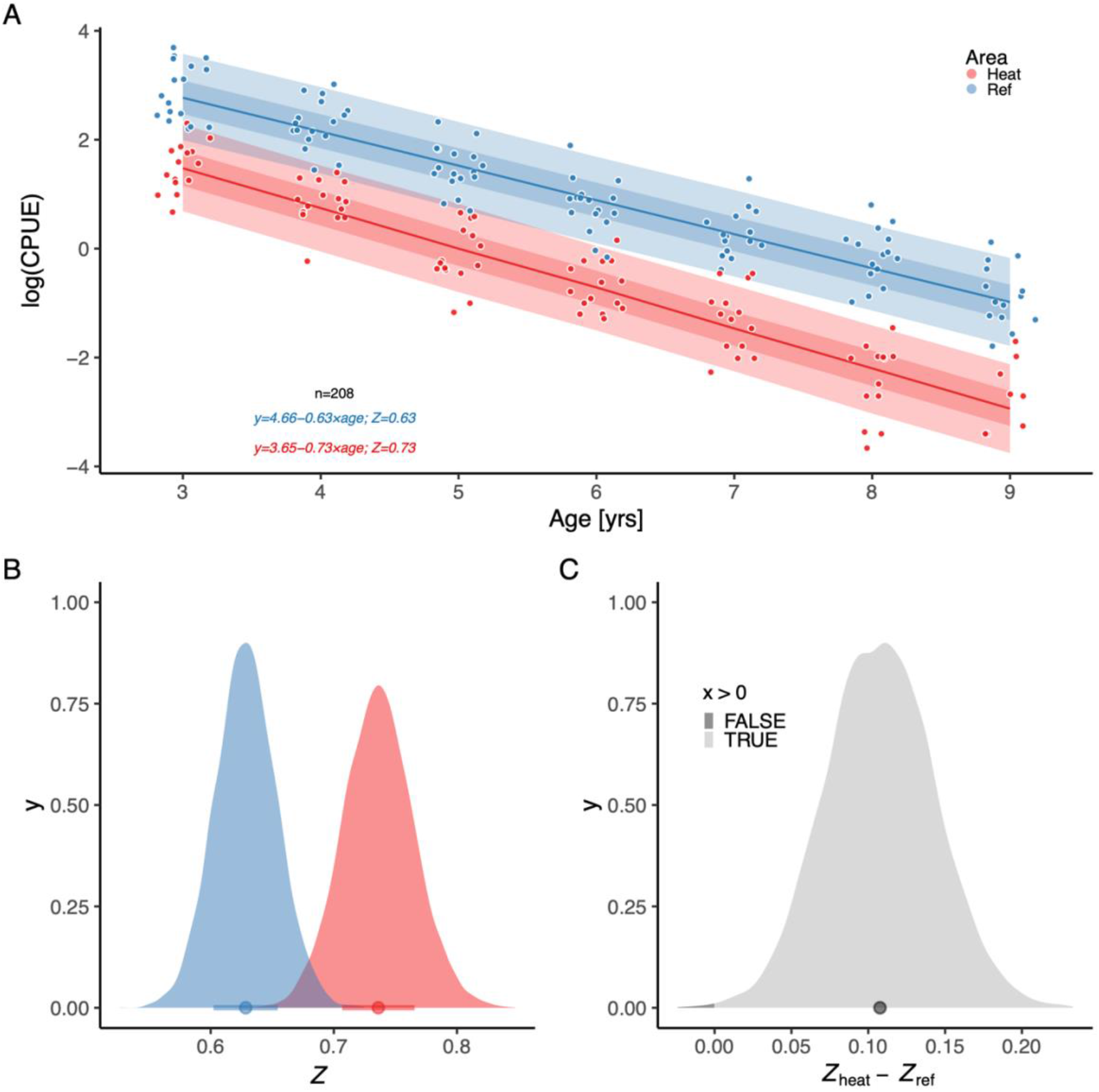
The instantaneous mortality rate (*Z*) is higher in the heated area (red) than in the reference (blue). Panel (A) shows the log(*CPUE*) as a function of *age*, where the slope corresponds to −*Z*. Lines show the posterior draws of the global posterior predictive distribution (without group-level effects) and the shaded areas correspond to the 50% and 90% credible intervals. The equation shows mean parameter estimates. Panel (B) shows the posterior distributions for mortality rate (*Z*_heat_ and *Z*_ref_), and (C) the distribution of their difference, where fill color depicts the area below 0 (0.07%).

Lastly, analysis of the size- and age-structure in the two areas revealed that, despite the faster growth rates, higher mortality and larger maximum sizes in the heated area for fish of all sizes, the size-spectrum exponents were largely similar. The size-spectrum exponent was only slightly larger in the heated area (Fig. 5A), and their 95% confidence intervals largely overlap (Fig. 5C). However, results from the lognormal model fitted to the size- and age-distributions revealed that the average size was two centimeters longer and the average age 0.4 years younger in the heated compared to the reference area (Fig. 6).

**Fig. 5.**
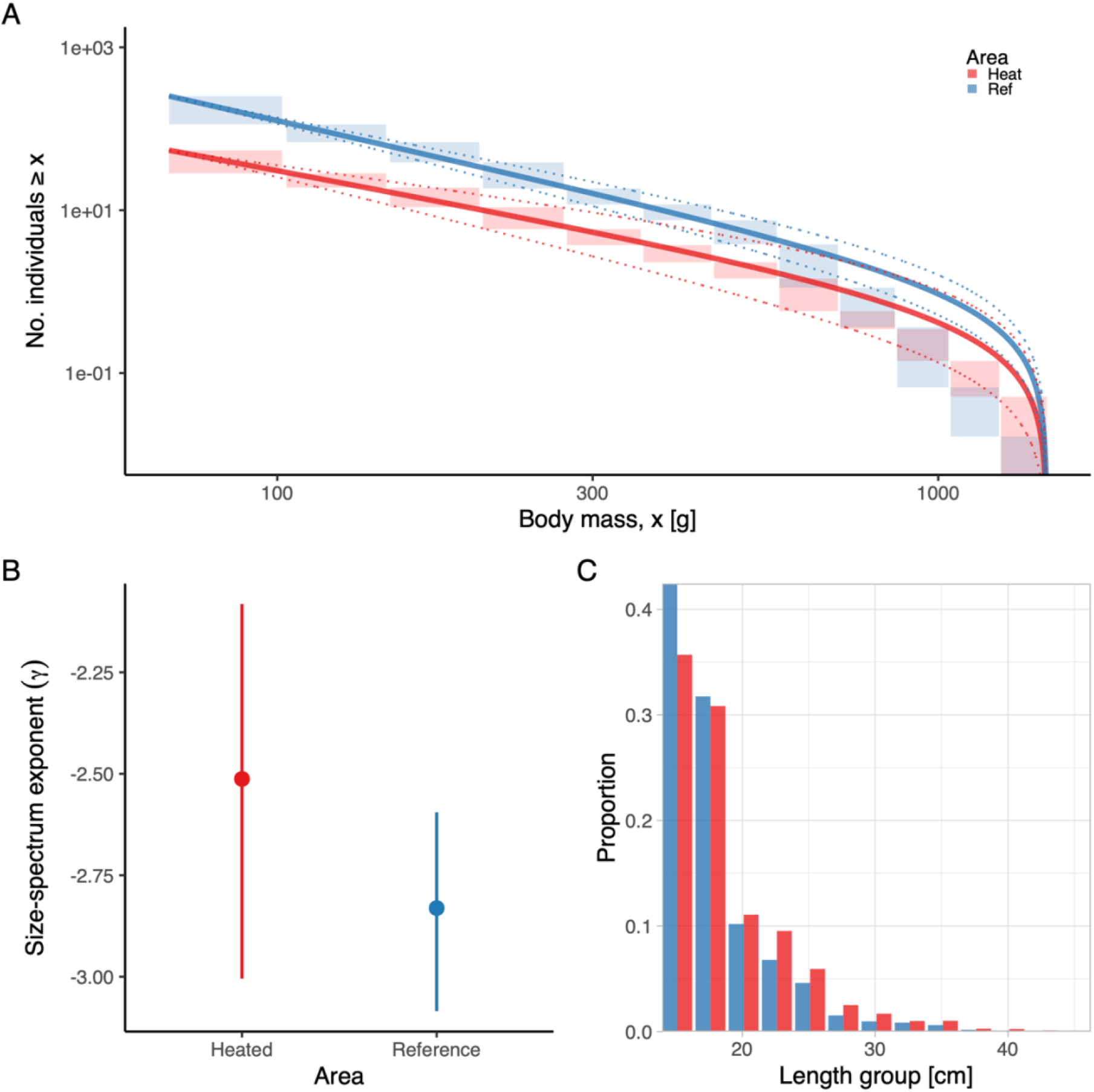
The heated area (red) has a larger proportion of large fish than the reference area (blue), illustrated both in terms of the biomass size-spectrum (A), and histograms of proportions (C), but the difference in the slope of the size-spectra between the areas is not statistically clear (B). Panel (A) shows the size distribution and MLEbins fit (red and blue solid curve for the heated and reference area, respectively) with 95% confidence intervals indicated by dashed lines. The vertical span of rectangles illustrates the possible range of the number of individuals with body mass ≥ the body mass of individuals in that bin. Panel (B) shows the estimate of the size-spectrum exponent, *γ*, and vertical lines depict the 95% confidence interval. Panel (C) illustrates histograms of length groups in the heated and reference area as proportions (for all years pooled).

**Fig. 6.**
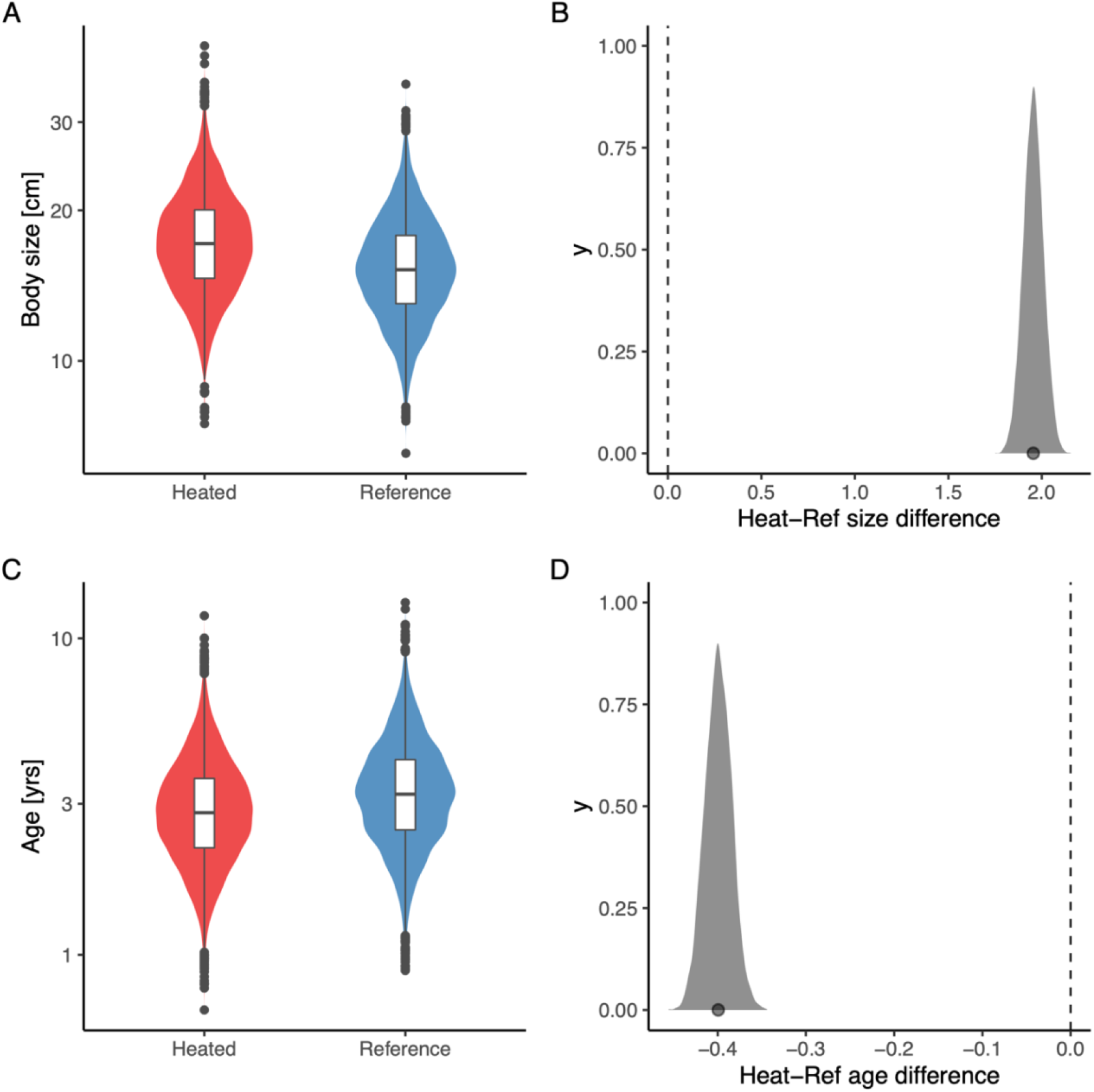
The average size is larger (A, B), but the average age (C, D) is younger in the heated area compared to the reference area. The violin plots (A, C) depict the posterior draws of the global posterior predictive distribution (without group-level effects) for mean size and age from the lognormal model, respectively, with the random year effect omitted, while the density plots (B, D) depict the difference between posterior draws for the area-parameters representing the means. The distributions in A, C, have higher variance than the posterior draws used to calculate densities in B, D, because they are posterior predictive distributions and thus also include residual error. The average size is 2 cm larger in the heated area, and the average age is 0.4 years younger (B, D).

## Discussion

Our study provides strong evidence for warming-induced differentiation in growth and mortality in a natural population of an unexploited, temperate fish species exposed to an ecosystem-scale experiment with 5–10 °C above normal temperatures for more than two decades. Interestingly, these effects largely, but not completely, counteract each other when it comes to population size-structure—while the fish are younger, they are also larger on average. However, differences in the rate of decline in abundance over size are less pronounced between the areas. It is difficult to generalize these findings since it is a study on only a single species. It is however, a unique climate change experiment, as experimental studies on fish to date are much shorter and often on scales much smaller than whole ecosystems, and long time series of biological samples exist mainly for commercially exploited fish species (Baudron et al., 2014; Smoliński et al., 2020; Thresher et al., 2007) (in which fisheries exploitation affects size-structure both directly and indirectly by selecting for fast growing individuals). While factors other than temperature could have contributed to the observed elevated growth and mortality, the temperature contrast is unusually large for natural systems (i.e., 5–10 °C, which can be compared to the 1.35 *°C* change in the Baltic Sea between 1982 and 2006 (Belkin, 2009)). Moreover, heating occurred at the scale of a whole ecosystem, which makes the findings highly relevant in the context of global warming.

Interestingly, our findings contrast with both broader predictions about declining mean or adult body sizes based on the GOLT hypothesis (Cheung et al., 2013; Pauly, 2021), and with intraspecific patterns such as the TSR (temperature-size rule, Atkinson (1994)). The contrasts lie in that both asymptotic size and size-at-age of mature individuals, as well as the proportion of larger individuals were slightly larger and higher in the heated area—despite the elevated mortality rates. This result was unexpected for two reasons: optimum growth temperatures generally decline with body size within species under food satiation in experimental studies (Lindmark et al., 2022), and fish tend to mature at smaller body size and allocate more energy into reproduction as it gets warmer (Niu et al., in press; Wootton et al., 2022). Both patterns have been used to explain how growth can increase for small and young fish, while large and old fish typically do not benefit from warming. Our study species is no exception to these rules (Huss et al., 2019; Karås and Thoresson, 1992; Niu et al., in press; Sandström et al., 1995). This suggests that growth dynamics under food satiation may not be directly proportional to those under natural feeding conditions (Railsback, 2022). It could also mean that while temperatures approach optimum for growth in the warmest months of the year (Huss et al., 2019), the exposure to such high temperatures is not enough to cause strong reductions in growth and eventually size-at-age. Moreover, our results suggest that growth changes emerge not only from direct physiological responses to increased temperatures, but also from warming-induced changes in the food web, e.g., prey productivity, diet composition and trophic transfer efficiencies (Gårdmark and Huss, 2020). It also highlights that we need to focus on understanding to what extent the commonly observed increase in size-at-age for juveniles in warm environments can be maintained as they grow older.

Our finding that mortality rates were higher in the heated area was expected—warming leads to faster metabolic rates (faster ‘pace of life’), which in turn is associated with shorter life span (Brown et al., 2004; McCoy and Gillooly, 2008; Munch and Salinas, 2009) (higher “physiological” mortality). Extreme temperatures, which may be more common in warmed systems under natural variability, can also be lethal if e.g., acute oxygen demands cannot be met (Sandblom et al., 2016). Warming may further increase predation mortality, as predators’ feeding rates increase in order to meet the higher demand of food (Biro et al., 2007; Pauly, 1980; Ursin, 1967). However, most evidence to date of the temperature dependence of mortality rates in natural populations stem from across species studies (Gislason et al., 2010; Pauly, 1980; Thorson et al., 2017) (but see (Berggren et al., 2021; Biro et al., 2007)). Across species relationships are not necessarily determined by the same processes as within species relationships; thus, our finding of warming-induced mortality in a heated vs control environment in two nearby con-specific populations is important.

Since a key question for understanding the implications of warming on ectotherm populations is if larger individuals in a population become rarer or smaller (Ohlberger, 2013; Ohlberger et al., 2018), within-species mortality and growth responses to warming need further study. Importantly, this requires accounting also for effects of warming on growth, and how responses in growth and mortality depend on each other. For instance, higher mortality (predation or natural, physiological mortality) can release intra-specific competition and thus increase growth. While e.g., benthic invertebrate density was not affected by the initial warming of the heated area (Sandström et al., 1995), warming-induced mortality may have led to higher benthic prey availability *per capita* for the studied perch. Conversely, altered growth and body sizes can lead to changes in size-specific mortality, such as predation or starvation, both which are expected to change with warming (Thunell, 2023). In conclusion, individual-level patterns such as the TSR may be of limited use for predicting changes on the population-level size structure as it does not concern changes in abundance-at-size via mortality. Mortality may, however, be an important driver of the observed shrinking of ectotherms (Peralta-Maraver and Rezende, 2021). Understanding the mechanisms by which the size- and age-distribution change with warming is critical for predicting how warming changes species functions and ecological roles (Audzijonyte et al., 2020; Fritschie and Olden, 2016; Gårdmark and Huss, 2020). Our findings demonstrate that a key to do this is to acknowledge temperature effects on both growth and mortality and how they interact.

## Acknowledgements

We thank all staff involved in data collection, Jens Olsson and Göran Sundblad for discussions, Christine Stawitz and an anonymous reviewer for feedback that greatly improved the manuscript. This study was supported by SLU Quantitative Fish and Fisheries Ecology.

## Code and Data Availability

All data and R code to reproduce the analyses can be downloaded from a GitHub repository (https://github.com/maxlindmark/warm-life-history) and will be archived on Zenodo upon publication. Researchers interested in using the data for purposes other than replicating our analyses are advised to request the data from the authors, as other useful information from the original data might not be included.

## Author Contributions

ML conceived the idea and designed the study and the statistical analysis. Data-processing, initial statistical analyses, and initial writing was done by MK and ML. AG contributed critically to all mentioned parts of the paper. All authors contributed to the manuscript writing and gave final approval for publication.

## References

Adill A, Mo K, Sevastik A, Olsson J, Bergström L. 2013. Biologisk recipientkontroll vid Forsmarks kärnkraftverk (in Swedish) (Rapport No. 2013:19). Öregrund.

Andersen KH. 2020. Size-based theory for fisheries advice. ICES J Mar Sci 77:2445–2455. doi:10.1093/icesjms/fsaa157

Andersen KH. 2019. Fish Ecology, Evolution, and Exploitation: A New Theoretical Synthesis. Princeton University Press.

Atkinson D. 1994. Temperature and organism size—A biological law for ectotherms? Advances in Ecological Research 25:1–58. doi:10.1016/S0065-2504(08)60212-3

Atkinson D, Sibly RM. 1997. Why are organisms usually bigger in colder environments? Making sense of a life history puzzle. Trends in Ecology & Evolution 12:235–239.

Audzijonyte A, Fulton E, Haddon M, Helidoniotis F, Hobday AJ, Kuparinen A, Morrongiello J, Smith AD, Upston J, Waples RS. 2016. Trends and management implications of human-influenced life-history changes in marine ectotherms. Fish and Fisheries 17:1005–1028. doi:10.1111/faf.12156

Audzijonyte A, Richards SA, Stuart-Smith RD, Pecl G, Edgar GJ, Barrett NS, Payne N, Blanchard JL. 2020. Fish body sizes change with temperature but not all species shrink with warming. Nat Ecol Evol 4:809–814. doi:10.1038/s41559-020-1171-0

Barneche DR, Jahn M, Seebacher F. 2019. Warming increases the cost of growth in a model vertebrate. Functional Ecology 33:1256–1266. doi:10.1111/1365-2435.13348

Barnett HK, Quinn TP, Bhuthimethee M, Winton JR. 2020. Increased prespawning mortality threatens an integrated natural-and hatchery-origin sockeye salmon population in the Lake Washington Basin. Fisheries Research 227:105527. doi:10.1016/j.fishres.2020.105527

Barnett LAK, Branch TA, Ranasinghe RA, Essington TE. 2017. Old-Growth Fishes Become Scarce under Fishing. Current Biology 27:2843–2848.e2. doi:10.1016/j.cub.2017.07.069

Baudron AR, Needle CL, Rijnsdorp AD, Marshall CT. 2014. Warming temperatures and smaller body sizes: synchronous changes in growth of North Sea fishes. Global Change Biology 20:1023–1031. doi:10.1111/gcb.12514

Belkin IM. 2009. Rapid warming of large marine ecosystems. Progress in Oceanography 81:207–213.

Berggren T, Bergström U, Sundblad G, Östman Ö. 2021. Warmer water increases early body growth of northern pike (Esox lucius) but mortality has larger impact on decreasing body sizes. Can J Fish Aquat Sci. doi:10.1139/cjfas-2020-0386

Bernhardt JR, Sunday JM, Thompson PL, O’Connor MI. 2018. Nonlinear averaging of thermal experience predicts population growth rates in a thermally variable environment. Proceedings of the Royal Society B: Biological Sciences 285:20181076. doi:10.1098/rspb.2018.1076

Beverton RJH, Holt SJ. 1957. On the Dynamics of Exploited Fish Populations. Fishery Investigations London Series 2, Volume 19.

Biro PA, Post JR, Booth DJ. 2007. Mechanisms for climate-induced mortality of fish populations in whole-lake experiments. Proceedings of the National Academy of Sciences 104:9715–9719. doi:10.1073/pnas.0701638104

Björklund M, Aho T, Behrmann-Godel J. 2015. Isolation over 35 years in a heated biotest basin causes selection on MHC class IIß genes in the European perch (Perca fluviatilis L.). Ecol Evol 5:1440–1455. doi:10.1002/ece3.1426

Blanchard JL, Dulvy N, Jennings S, Ellis J, Pinnegar J, Tidd A, Kell L. 2005. Do climate and fishing influence size-based indicators of Celtic Sea fish community structure? ICES Journal of Marine Science 62:405–411. doi:10.1016/j.icesjms.2005.01.006

Blanchard JL, Heneghan RF, Everett JD, Trebilco R, Richardson AJ. 2017. From bacteria to whales: Using functional size spectra to model marine ecosystems. Trends in Ecology & Evolution 32:174–186. doi:10.1016/j.tree.2016.12.003

Brown JH, Gillooly JF, Allen AP, Savage VM, West GB. 2004. Toward a metabolic theory of ecology. Ecology 85:1771–1789. doi:10.1890/03-9000

Bürkner P-C. 2018. Advanced Bayesian Multilevel Modeling with the R Package brms. The R Journal 10:395–411.

Bürkner P-C. 2017. **brms**: An *R* Package for Bayesian Multilevel Models Using *Stan*. Journal of Statistical Software 80. doi:10.18637/jss.v080.i01

Cheung WWL, Sarmiento JL, Dunne J, Frölicher TL, Lam VWY, Deng Palomares ML, Watson R, Pauly D. 2013. Shrinking of fishes exacerbates impacts of global ocean changes on marine ecosystems. Nature Climate Change 3:254–258. doi:10.1038/nclimate1691

Dunn A, Francis RICC, Doonan IJ. 2002. Comparison of the Chapman–Robson and regression estimators of Z from catch-curve data when non-sampling stochastic error is present. Fisheries Research 59:149–159. doi:10.1016/S0165-7836(01)00407-6

Edwards A. 2020. sizeSpectra: Fitting Size Spectra to Ecological Data Using Maximum Likelihood.

Edwards AM, Robinson JPW, Blanchard JL, Baum JK, Plank MJ. 2020. Accounting for the bin structure of data removes bias when fitting size spectra. Marine Ecology Progress Series 636:19–33. doi:10.3354/meps13230

Edwards AM, Robinson JPW, Plank MJ, Baum JK, Blanchard JL. 2017. Testing and recommending methods for fitting size spectra to data. Methods in Ecology and Evolution 8:57–67. doi:10.1111/2041-210X.12641

Essington TE, Matta ME, Black BA, Helser TE, Spencer PD. 2022. Fitting growth models to otolith increments to reveal time-varying growth. Can J Fish Aquat Sci 79:159–167. doi:10.1139/cjfas-2021-0046

Fritschie KJ, Olden JD. 2016. Disentangling the influences of mean body size and size structure on ecosystem functioning: an example of nutrient recycling by a non-native crayfish. Ecology and Evolution 6:159–169. doi:10.1002/ece3.1852

Gabry J, Simpson D, Vehtari A, Betancourt M, Gelman A. 2019. Visualization in Bayesian workflow. J R Stat Soc A 182:389–402. doi:10.1111/rssa.12378

Gårdmark A, Huss M. 2020. Individual variation and interactions explain food web responses to global warming. Philosophical Transactions of the Royal Society B: Biological Sciences 375:20190449. doi:10.1098/rstb.2019.0449

Gardner JL, Peters A, Kearney MR, Joseph L, Heinsohn R. 2011. Declining body size: a third universal response to warming? Trends in Ecology & Evolution 26:285–291. doi:10.1016/j.tree.2011.03.005

Gelman A, Carlin J, Stern H, Rubin D. 2003. Bayesian Data Analysis. 2nd edition. Boca Raton: Chapman and Hall/CRC.

Gesmann M, Morris J. 2020. Hierarchical compartmental reserving modelsCasualty Actuarial Society. p. 4.

Gislason H, Daan N, Rice JC, Pope JG. 2010. Size, growth, temperature and the natural mortality of marine fish: Natural mortality and size. Fish and Fisheries 11:149–158. doi:10.1111/j.1467-2979.2009.00350.x

Heneghan RF, Hatton IA, Galbraith ED. 2019. Climate change impacts on marine ecosystems through the lens of the size spectrum. Emerging Topics in Life Sciences 3:233–243. doi:10.1042/ETLS20190042

Huss M, Lindmark M, Jacobson P, van Dorst RM, Gårdmark A. 2019. Experimental evidence of gradual size-dependent shifts in body size and growth of fish in response to warming. Glob Change Biol 25:2285–2295. doi:10.1111/gcb.14637

Karås P, Thoresson G. 1992. An application of a bioenergetics model to Eurasian perch (Perca fluviatilis L.). Journal of Fish Biology 41:217–230.

Kay M. 2019. tidybayes: Tidy Data and Geoms for Bayesian Models.

Lindmark M, Ohlberger J, Gårdmark A. 2022. Optimum growth temperature declines with body size within fish species. Global Change Biology 28:2259–2271. doi:10.1111/gcb.16067

Lorenzen K. 2016. Toward a new paradigm for growth modeling in fisheries stock assessments: Embracing plasticity and its consequences. Fisheries Research, Growth: theory, estimation, and application in fishery stock assessment models 180:4–22. doi:10.1016/j.fishres.2016.01.006

McCoy MW, Gillooly JF. 2008. Predicting natural mortality rates of plants and animals. Ecology Letters 11:710–716. doi:10.1111/j.1461-0248.2008.01190.x

Morrongiello JR, Thresher RE. 2015. A statistical framework to explore ontogenetic growth variation among individuals and populations: a marine fish example. Ecological Monographs 85:93–115. doi:10.1890/13-2355.1

Munch SB, Salinas S. 2009. Latitudinal variation in lifespan within species is explained by the metabolic theory of ecology. Proceedings of the National Academy of Sciences 106:13860–13864. doi:10.1073/pnas.0900300106

Niu J, Huss M, Vasemägi A, Gårdmark A. in press. Decades of warming alters maturation and reproductive investment in fish. Ecosphere.

Ohlberger J. 2013. Climate warming and ectotherm body size – from individual physiology to community ecology. Functional Ecology 27:991–1001. doi:10.1111/1365-2435.12098

Ohlberger J, Ward EJ, Schindler DE, Lewis B. 2018. Demographic changes in Chinook salmon across the Northeast Pacific Ocean. Fish and Fisheries 19:533–546. doi:10.1111/faf.12272

Oke KB, Mueter FJ, Litzow MA. 2022. Warming leads to opposite patterns in weight-at-age for young versus old age classes of Bering Sea walleye pollock. Can J Fish Aquat Sci. doi:10.1139/cjfas-2021-0315

Pauly D. 2021. The gill-oxygen limitation theory (GOLT) and its critics. Science Advances 7:eabc6050. doi:10.1126/sciadv.abc6050

Pauly D. 1980. On the interrelationships between natural mortality, growth parameters, and mean environmental temperature in 175 fish stocks. ICES Journal of Marine Science 39:175–192. doi:10.1093/icesjms/39.2.175

Peralta-Maraver I, Rezende EL. 2021. Heat tolerance in ectotherms scales predictably with body size. Nat Clim Chang 11:58–63. doi:10.1038/s41558-020-00938-y

R Core Team. 2020. R: A Language and Environment for Statistical Computing. R Foundation for Statistical Computing. Vienna, Austria.

Railsback SF. 2022. What We Don’t Know About the Effects of Temperature on Salmonid Growth. Transactions of the American Fisheries Society 151:3–12. doi:10.1002/tafs.10338

Rindorf A, Jensen H, Schrum C. 2008. Growth, temperature, and density relationships of North Sea cod (Gadus morhua). Canadian Journal of Fisheries and Aquatic Sciences 65:456–470. doi:10.1139/f07-150

Sandblom E, Clark TD, Gräns A, Ekström A, Brijs J, Sundström LF, Odelström A, Adill A, Aho T, Jutfelt F. 2016. Physiological constraints to climate warming in fish follow principles of plastic floors and concrete ceilings. Nat Commun 7:11447. doi:10.1038/ncomms11447

Sandström O, Neuman E, Thoresson G. 1995. Effects of temperature on life history variables in perch. Journal of Fish Biology 47:652–670. doi:10.1111/j.1095-8649.1995.tb01932.x

Sheldon RW, Sutcliffe WH, Prakash A. 1973. The Production of Particles in the Surface Waters of the Ocean with Particular Reference to the Sargasso Sea1. Limnology and Oceanography 18:719–733. doi:10.4319/lo.1973.18.5.0719

Sheridan JA, Bickford D. 2011. Shrinking body size as an ecological response to climate change. Nature Climate Change 1:401–406. doi:10.1038/nclimate1259

Smoliński S, Deplanque-Lasserre J, Hjörleifsson E, Geffen AJ, Godiksen JA, Campana SE. 2020. Century-long cod otolith biochronology reveals individual growth plasticity in response to temperature. Sci Rep 10:16708. doi:10.1038/s41598-020-73652-6

Thoresson G. 1996. Metoder för övervakning av kustfiskbestånd (in Swedish) (No. 3), Kustrapport. Öregrund: Kustlaboratoriet, Fiskeriverket.

Thorson JT, Munch SB, Cope JM, Gao J. 2017. Predicting life history parameters for all fishes worldwide. Ecological Applications 27:2262–2276. doi:10.1002/eap.1606

Thresher RE, Koslow JA, Morison AK, Smith DC. 2007. Depth-mediated reversal of the effects of climate change on long-term growth rates of exploited marine fish. Proceedings of the National Academy of Sciences, USA 104:7461–7465.

Thunell V. 2023. Fish life histories in a warming climate – a mechanistic basis for change and a community context. (PhD Thesis). Uppsala: Swedish University of Agricultural Sciences.

Ursin E. 1967. A Mathematical Model of Some Aspects of Fish Growth, Respiration, and Mortality. Journal of the Fisheries Research Board of Canada 24:2355–2453. doi:10.1139/f67-190

van Gemert R, Andersen KH. 2018. Implications of late-in-life density-dependent growth for fishery size-at-entry leading to maximum sustainable yield. ICES Journal of Marine Science 75:1296–1305. doi:10.1093/icesjms/fsx236

Vehtari A, Gabry J, Magnusson M, Yao Y, Bürkner P, Paananen T, Gelman A. 2020. loo: Efficient leave-one-out cross-validation and WAIC for Bayesian models.

Vehtari A, Gelman A, Gabry J. 2017. Practical Bayesian model evaluation using leave-one-out cross-validation and WAIC. Stat Comput 27:1413–1432. doi:10.1007/s11222016-9696-4

von Bertalanffy L. 1938. A quantitative theory of organic growth (inquiries on growth laws. II). Human Biology 10:181–213.

Wesner JS, Pomeranz JPF. 2021. Choosing priors in Bayesian ecological models by simulating from the prior predictive distribution. Ecosphere 12:e03739. doi:10.1002/ecs2.3739

White EP, Ernest SKM, Kerkhoff AJ, Enquist BJ. 2007. Relationships between body size and abundance in ecology. Trends in Ecology & Evolution 22:323–330.

Wickham H, Averick M, Bryan J, Chang W, D’Agostino McGowan L, François R, Grolemund G, Alex H. 2019. Welcome to the tidyverse. Journal of Open Source Software 1686. doi:https://doi.org/10.21105/joss.01686

Wilson EO. 1992. The Diversity of Life. Cambridge: Harvard University Press.

Wootton HF, Morrongiello JR, Schmitt T, Audzijonyte A. 2022. Smaller adult fish size in warmer water is not explained by elevated metabolism. Ecology Letters 25:1177–1188. doi:10.1111/ele.13989

Yvon-Durocher G, Montoya JM, Trimmer M, Woodward G. 2011. Warming alters the size spectrum and shifts the distribution of biomass in freshwater ecosystems. Global Change Biology 17:1681–1694. doi:10.1111/j.1365-2486.2010.02321.x

