## Supporting Information for "Larger but younger fish when growth outpaces mortality in heated ecosystem"

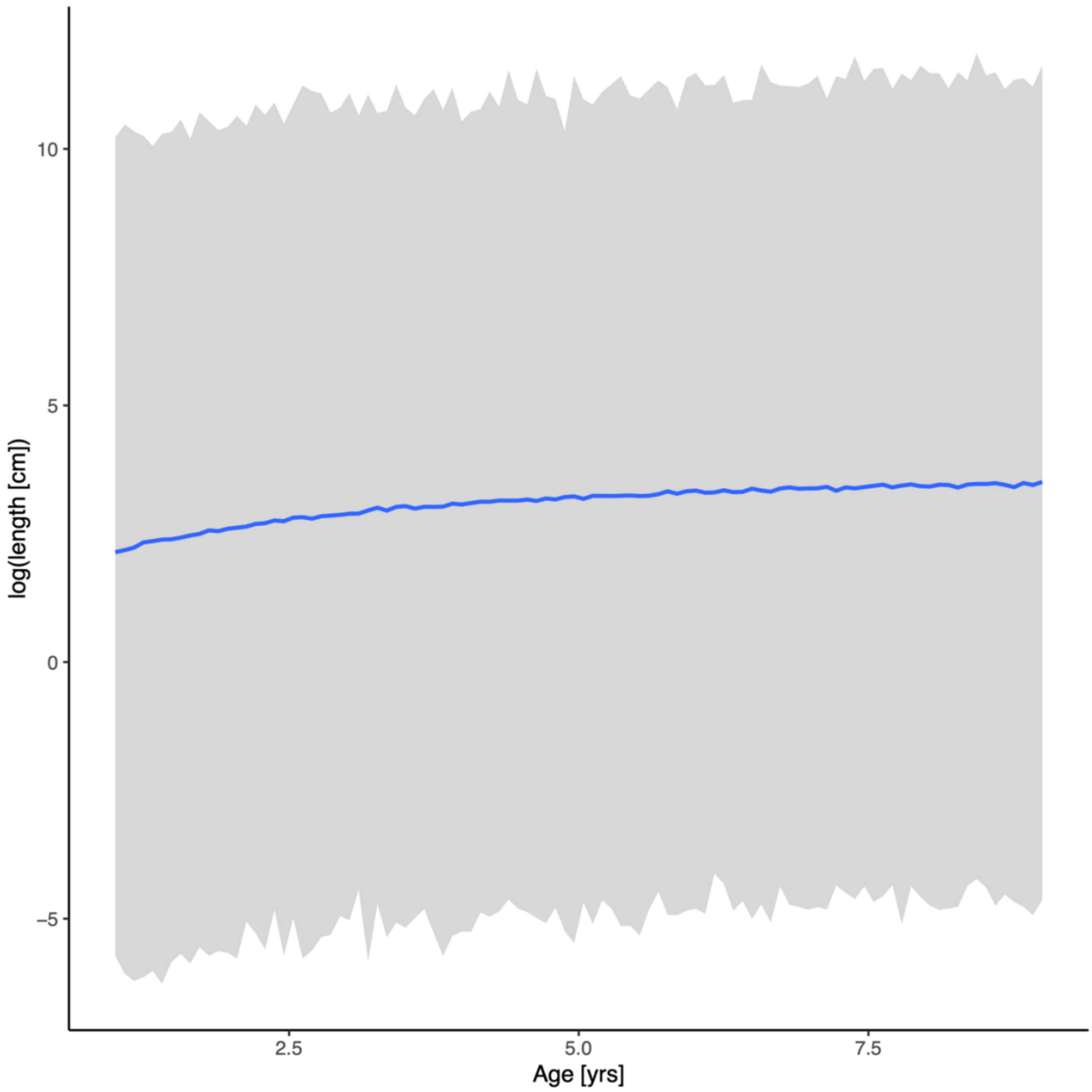


**Fig. S1.** Prior predictive distribution for the von Bertalanffy growth equation (posterior draws from the prior only, ignoring the likelihood). The solid line is the median and the shaded area is the 95% credible intervals.

**Table S1.** Comparison of von Bertalanffy growth models with different combinations of shared and area-specific parameters (ordered by difference in expected log pointwise density (elpd) from the best model). Note that in all models, $L_{\infty j}$ and $K_j$ vary among cohorts.

| **Model Name** | **Model structure** | **elpd_diff** |
|---|---|---|
| M1 | Area-specific $L_{\infty j}$, $K_j$ and $t_0$ | 0 |
| M4 | Area-specific $L_{\infty j}$ and $K_j$, common $t_0$ | -9.8 |
| M2 | Area-specific $K_j$, common $t_0$ and $L_{\infty j}$ | -112 |
| M3 | Area-specific $t_0$ and $L_{\infty j}$, common $K_j$ | -152 |
| M7 | Area-specific $L_{\infty j}$, common $K_j$ and $t_0$ | -158 |
| M6 | Area-specific $K_j$, common $t_0$ and $L_{\infty j}$ | -175 |
| M5* | Area-specific $t_0$, common $K_j$ and $L_{\infty j}$ | -1338 |
| M8* | Common $t_0$, $K_j$ and $L_{\infty j}$ | -2155 |

* Models did not converge

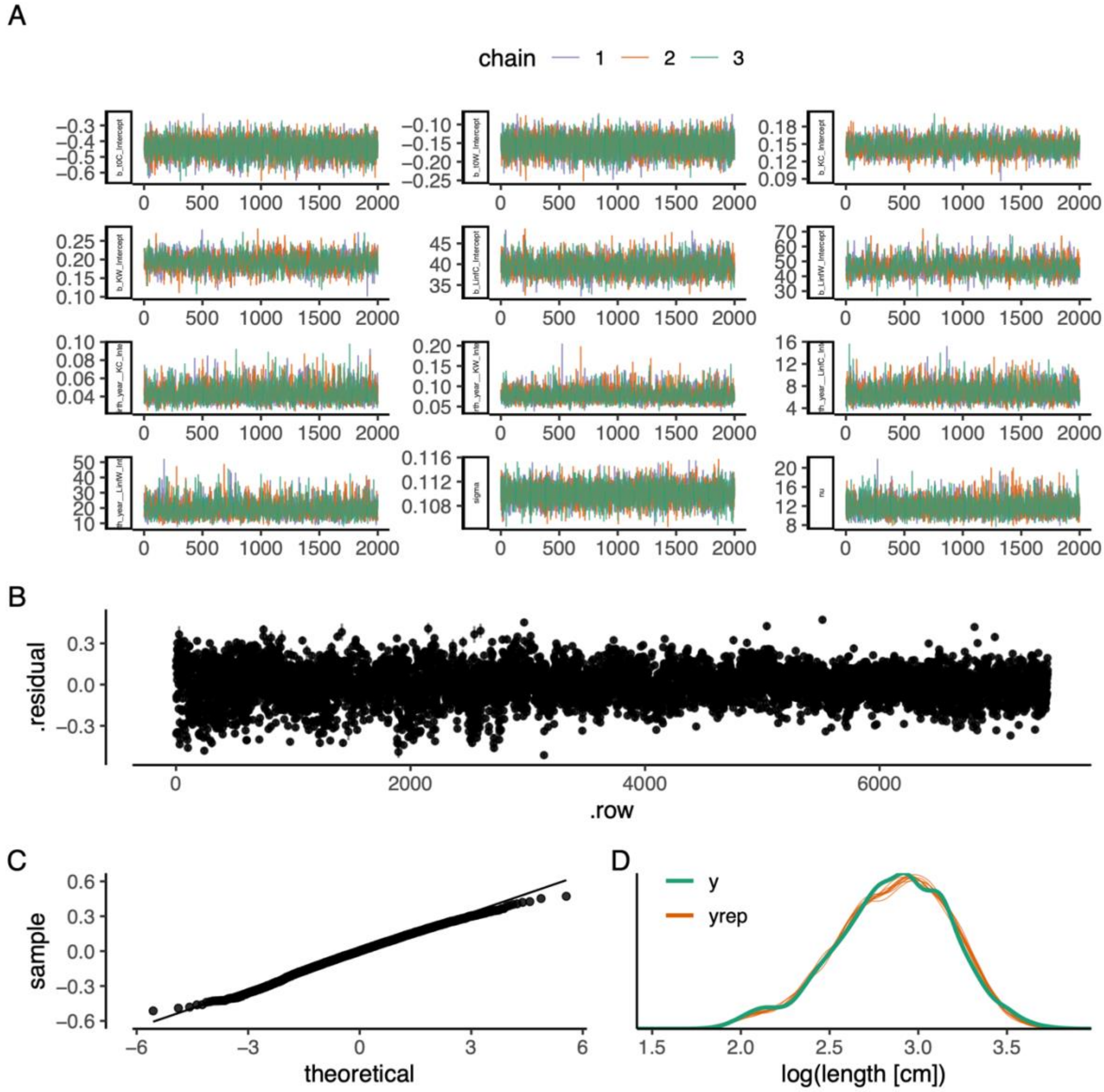


**Fig. S2.** The best model of the von Bertalanffy growth equation: (A) traceplot to illustrate chain convergence for key (population-level) parameters, (B) residuals, (C) QQ-plot and (D) posterior predictive check (D).

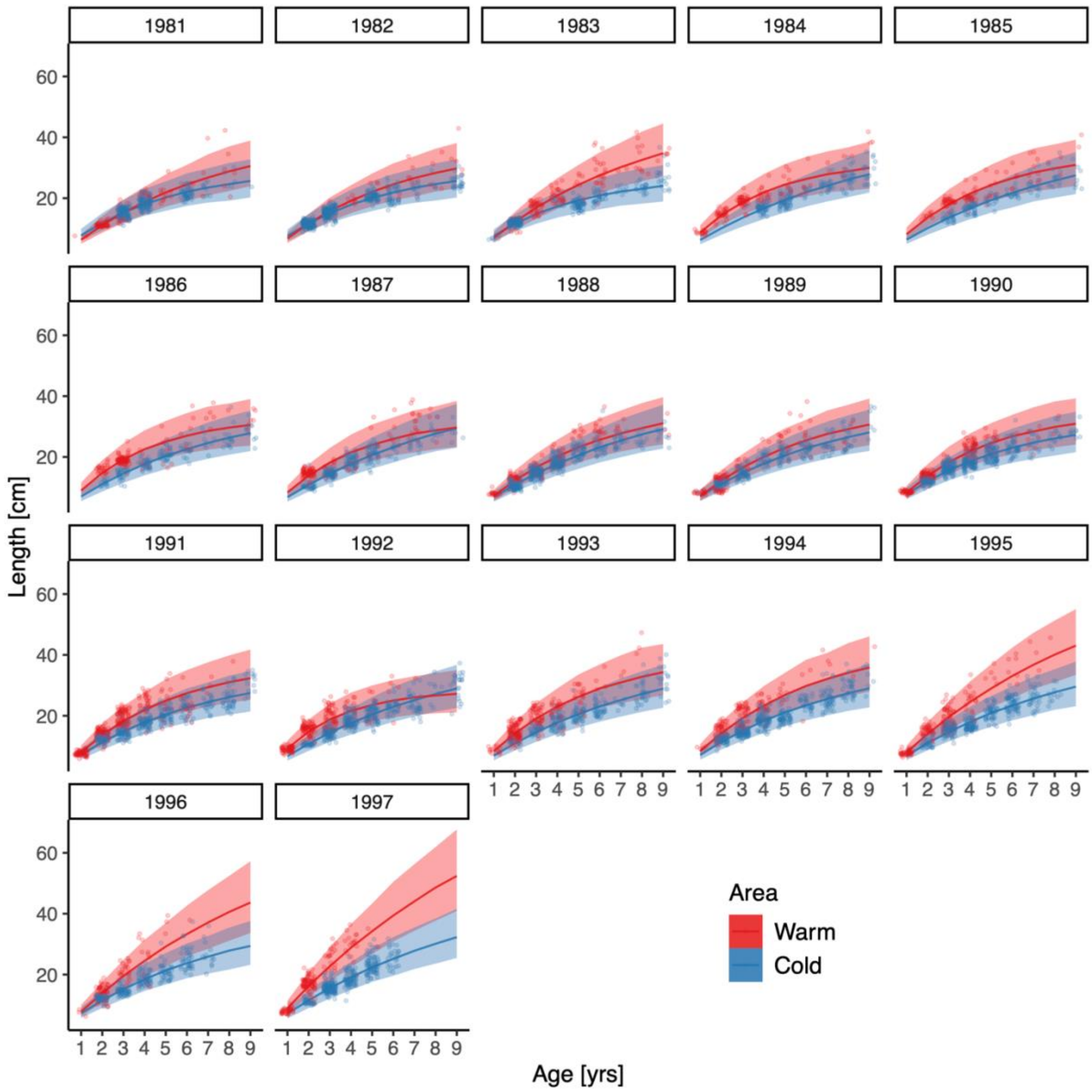


**Fig. S3.** Cohort-specific predictions from the best von Bertalanffy model (i.e., with cohort-varying $L_\infty$ and $K$). Points correspond to data; solid lines correspond to the median of the posterior prediction from the model and the shaded area corresponds to the 95% credible interval.

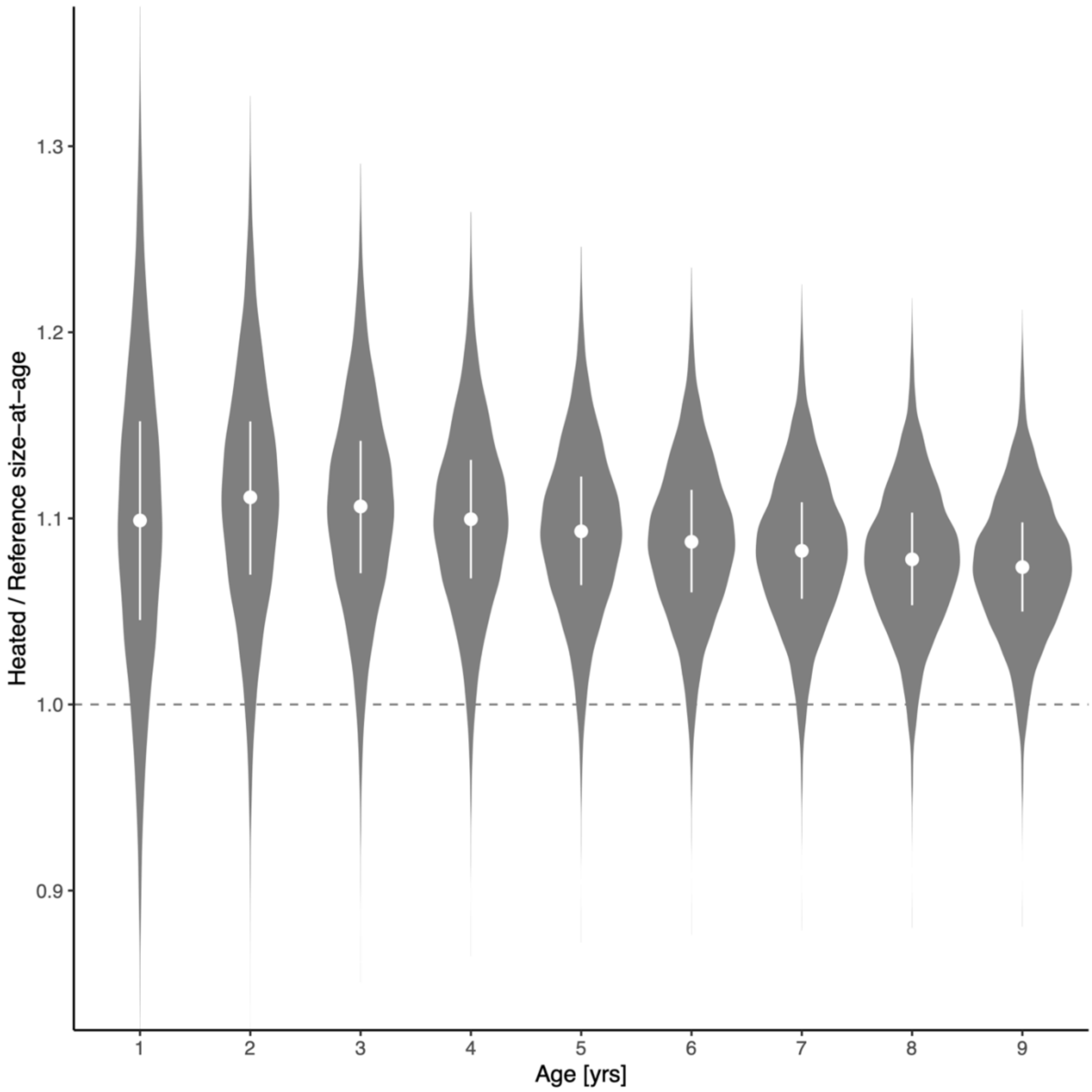


**Fig. S4.** The average length-at-age is larger for fish of all ages in the heated enclosed bay compared to the reference area, and the relative difference declines very slightly with age. Violin plots depict size-at-age in the heated relative to the reference area, based on draws from expectation of the posterior predictive distribution (without random effects). The points and vertical lines depict the median and the interquartile range.

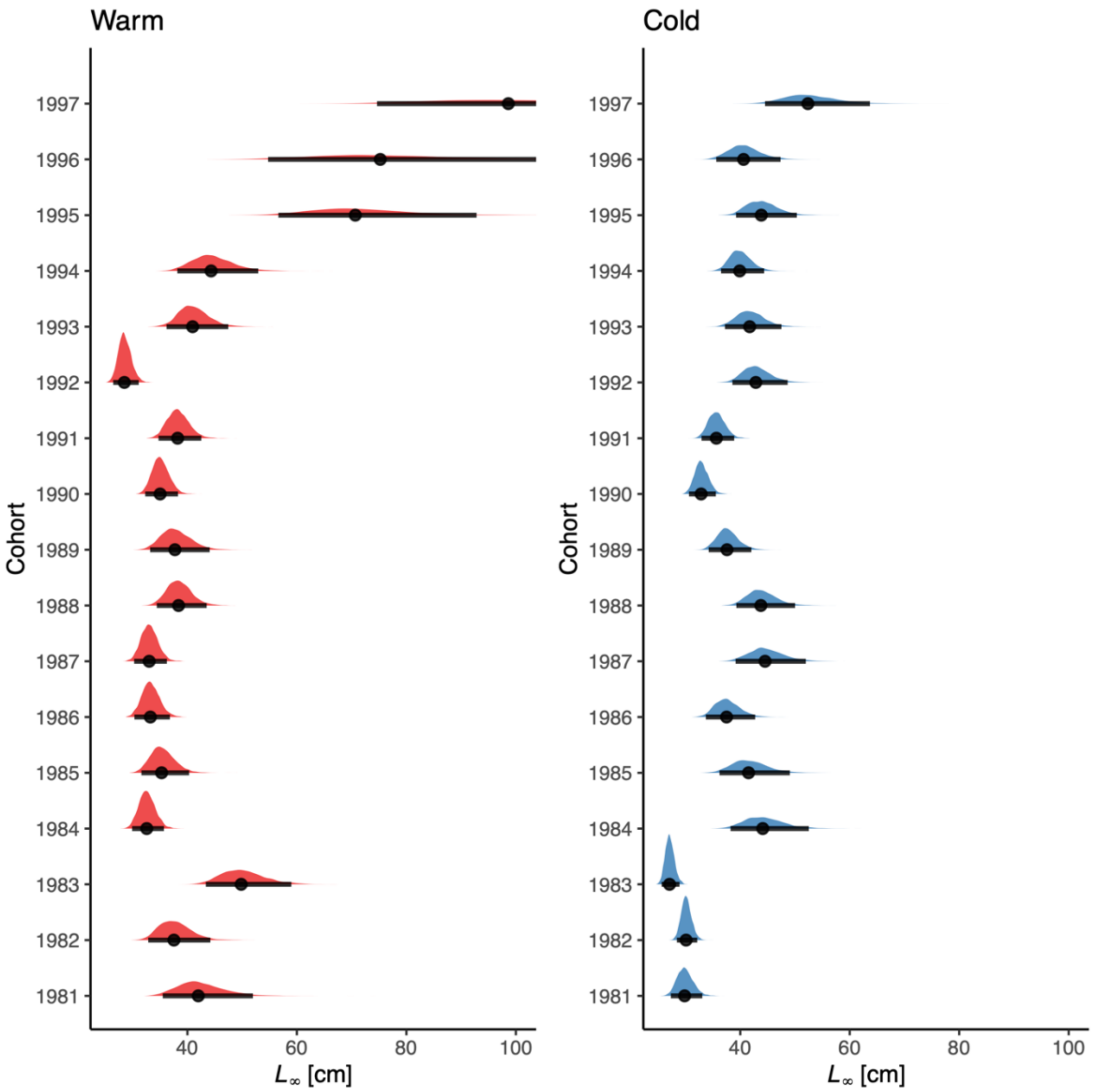


**Fig. S5.** Posterior distributions of the cohort-varying $L_\infty$ parameter in the best von Bertalanffy growth model. Points correspond to the median and the horizontal lines correspond to the 95% credible interval. Note that the distributions of $L_\infty$ in the warm areas extend beyond the x-axis for cohorts 1995–1997 (also evident in Fig. S3). The range of the x-axis was set to be wide enough to include the posterior medians of the larger estimates but narrow enough to allow for comparison between the other cohorts and areas.

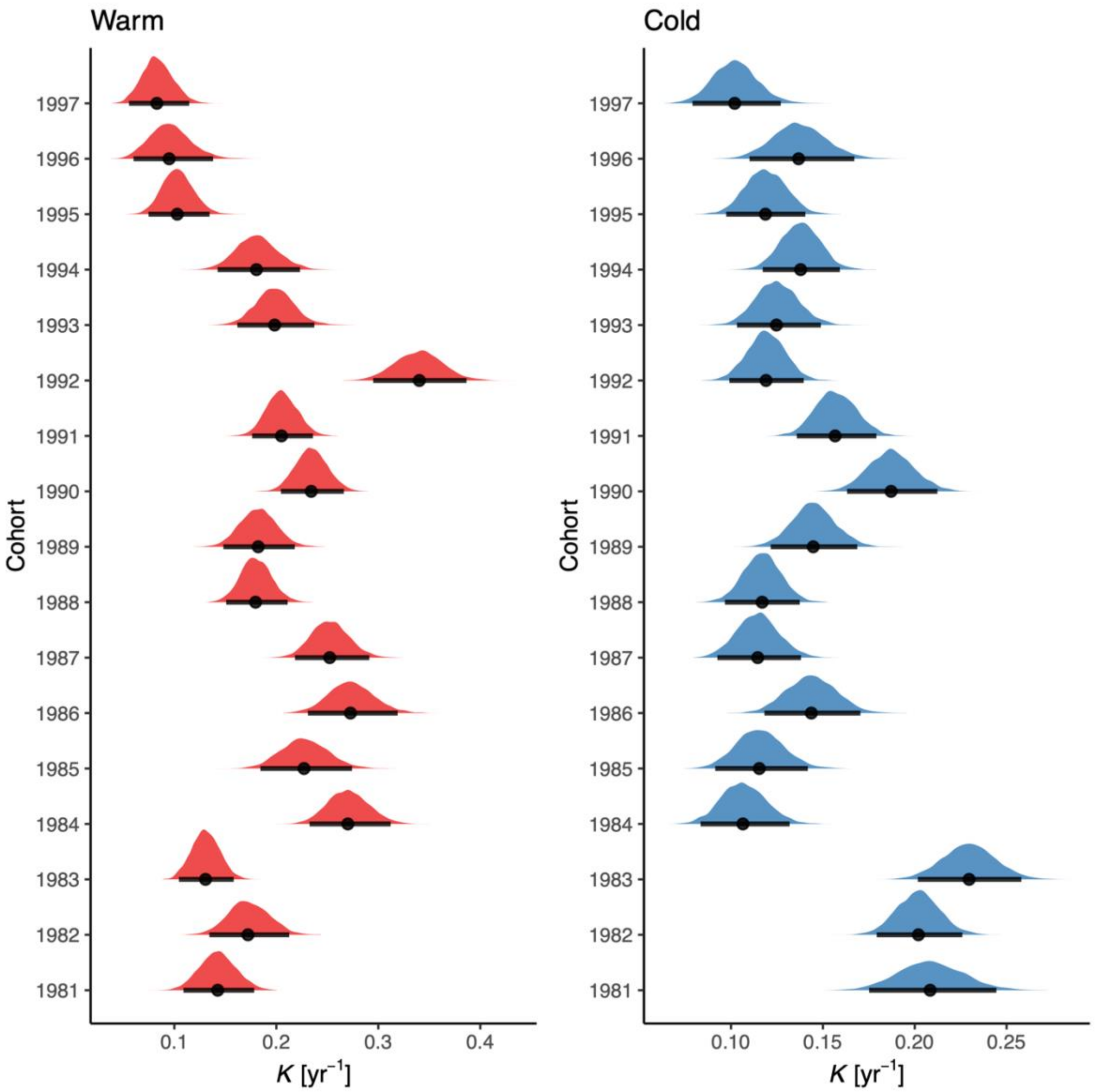


**Fig. S6.** Posterior distributions of the cohort-varying *K* parameter in the von Bertalanffy model. Points correspond to the median and the horizontal lines correspond to the 95% credible interval.

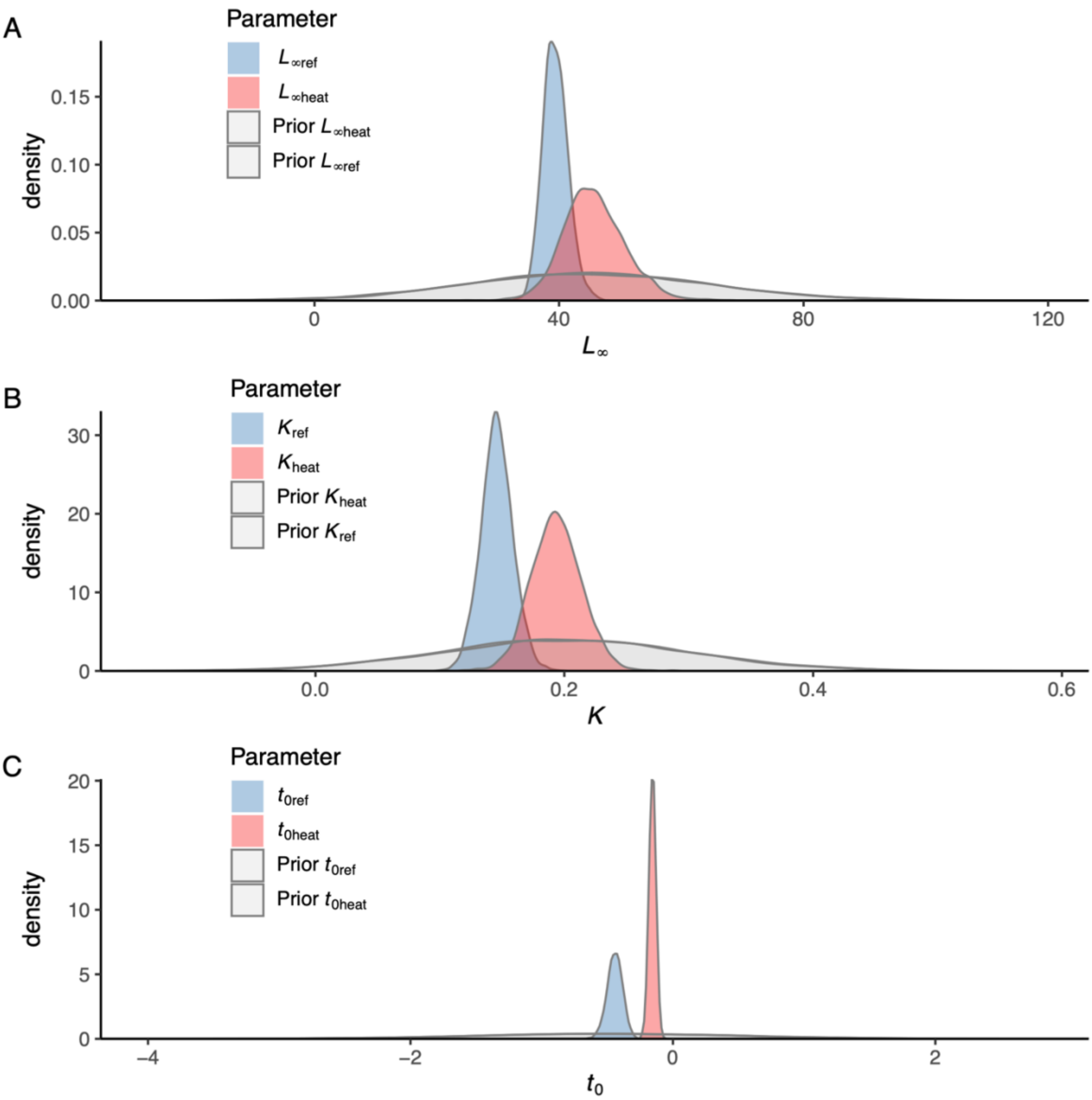


**Fig. S7.** Prior vs posterior distributions for parameters $L_{\infty}$ (A), $K$ (B) and $t_0$ (C) in the best model of the von Bertalanffy growth equation.

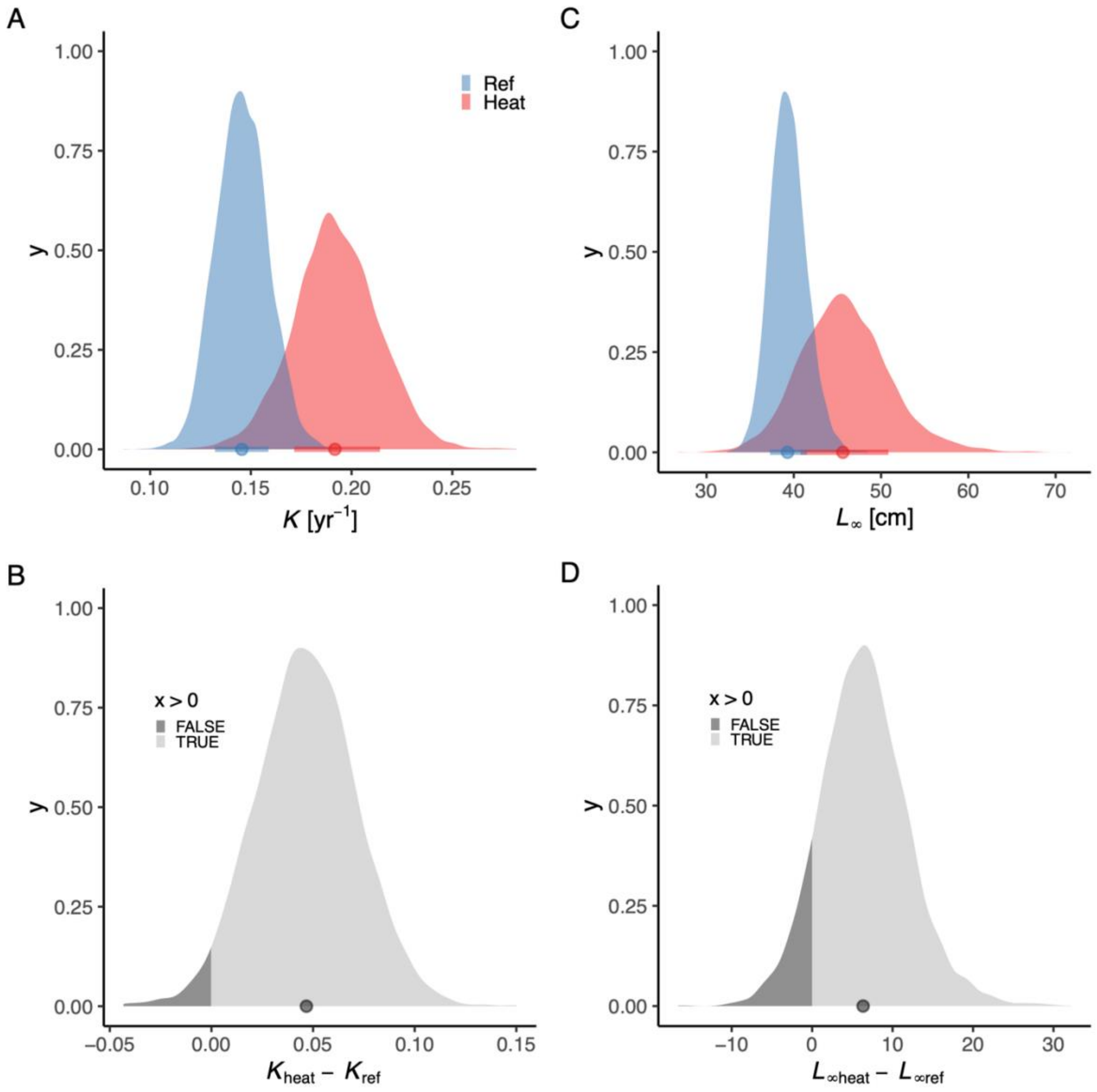


**Fig. S8.** Panel (A) depicts the posterior distributions for Brody growth coefficient (parameters $K_{\mathrm{heat}}$ (red) and $K_{\mathrm{ref}}$ (blue)) and (B) the distribution of their difference. Panel (C) depicts the posterior distributions for asymptotic length (parameters $L_{\infty\mathrm{heat}}$ and $L_{\infty\mathrm{ref}}$), and (D) the distribution of their difference (3%). The fill color depicts the area below 0 (3% and 11% for $K$ and $L_{\infty}$, respectively).

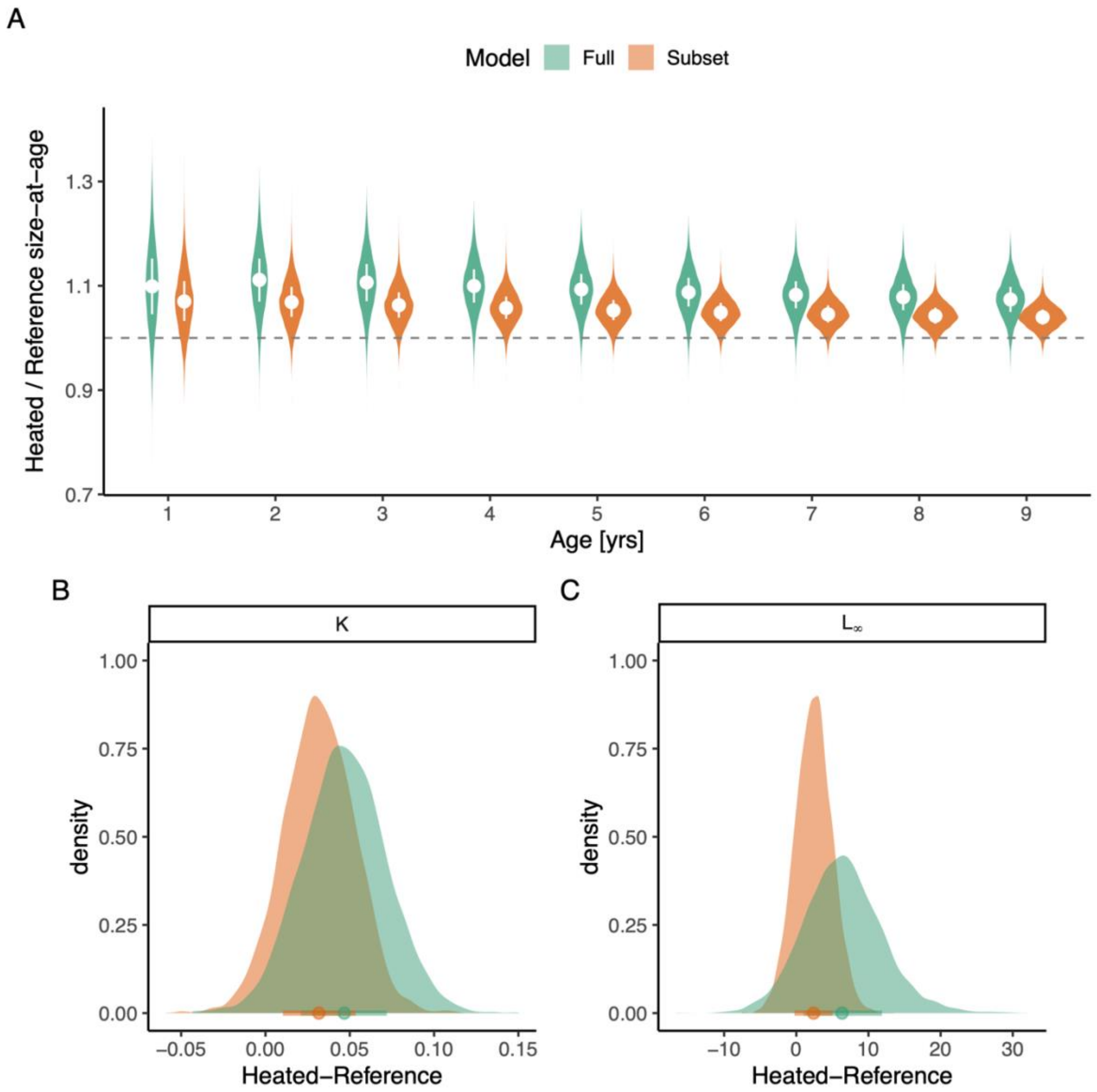


**Fig. S9.** Analysis of sensitivity of including the most recent cohorts, with a smaller age-range and therefore less certain estimates of $L_\infty$. Panel (A) depicts predicted size-at-age from the full model (green) and the same model fitted without cohorts 1995-1997. The violin plots depict size-at-age in the heated relative to the reference area, based on draws from expectation of the posterior predictive distribution (without random effects). The points and vertical lines depict the median and the interquartile range. Panels (B) and (C) depict the posterior distribution of differences in $K$ and $L_\infty$, respectively, whehere color again indicates full or subset models.

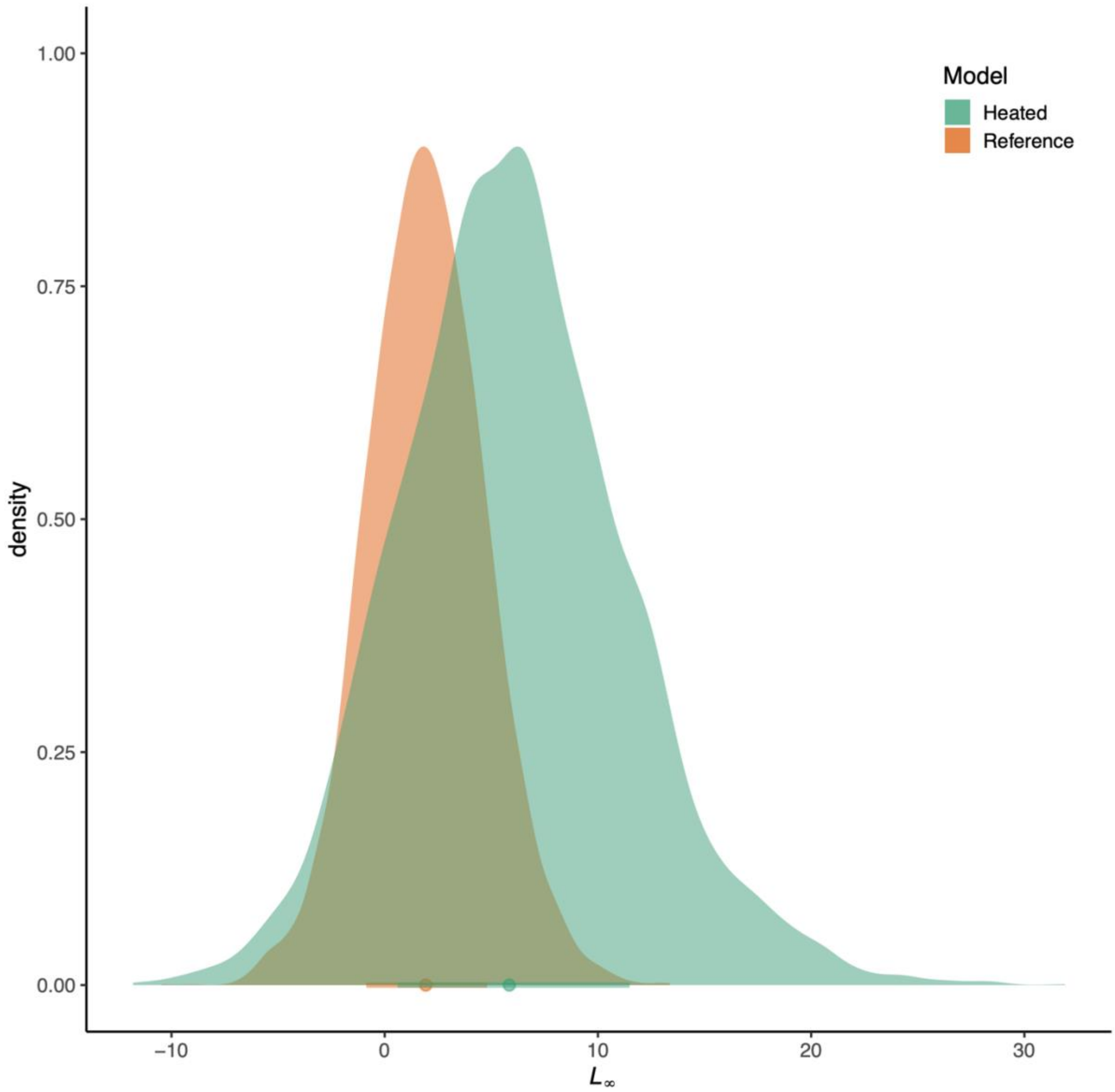


**Fig. S10.** Analysis of sensitivity of including the most recent cohorts, with a smaller age-range and therefore less certain estimates of $L_\infty$. The distributions depict the differences between the full and the subset models' posterior distribution for $L_\infty$, with colors corresponding to the estimate for the heated and reference area.

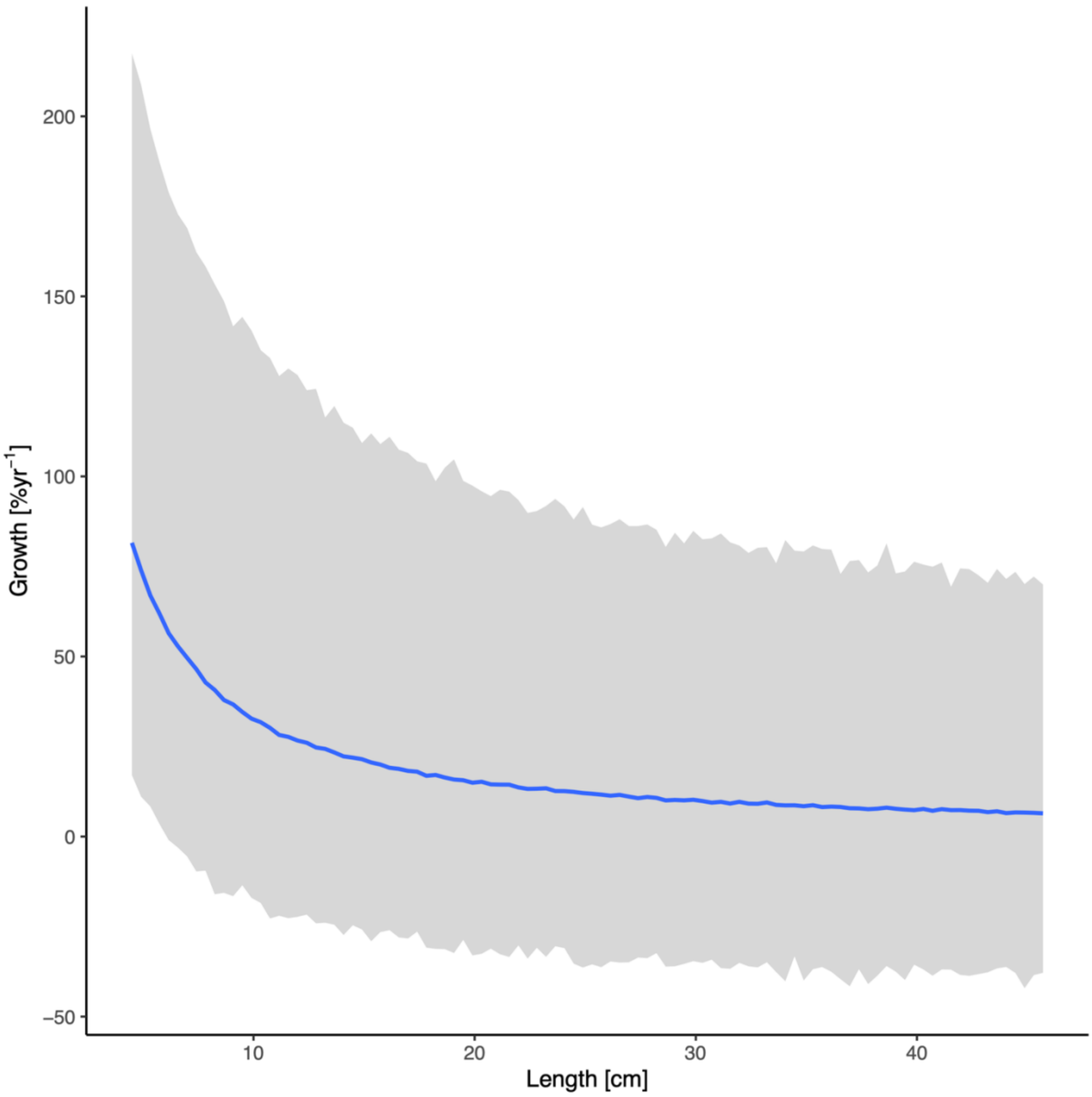


**Fig. S11.** Prior predictive distribution for the allometric growth model (posterior draws from the prior only, ignoring the likelihood). The solid line is the median and the shaded area is the 95% credible intervals.

**Table S2.** Comparison of allometric growth models with common or unique $\theta$-parameter (exponent in the allometric growth model), ordered by difference in expected log pointwise density (elpd) from the best model.

| **Model Name** | **Model structure** | **elpd_diff** |
|---|---|---|
| M1 | Intercept ($\alpha_{j[i],k[i]}$) varying across individuals within cohorts, fixed, area-specific slope ($\theta_{ref}, \theta_{heat}$) | 0 |
| M2 | Intercept ($\alpha_{j[i],k[i]}$) varying across individuals within cohorts, “fixed” common slope ($\theta$) | -1.9 |

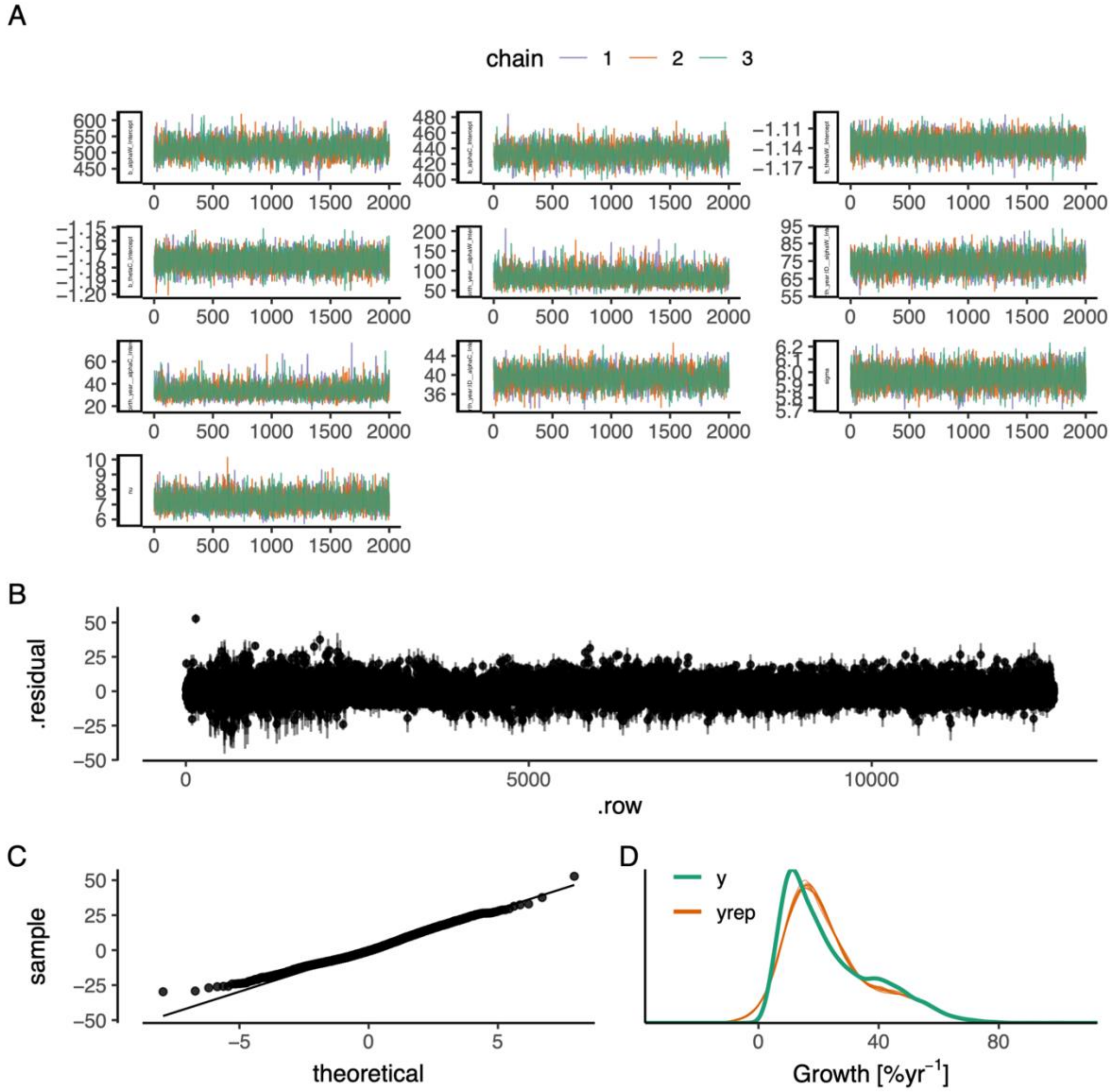


**Fig. S12.** The best allometric growth model: (A) traceplot to illustrate chain convergence for key (population-level) parameters, (B) residuals, (C) QQ-plot and (D) posterior predictive check (D).

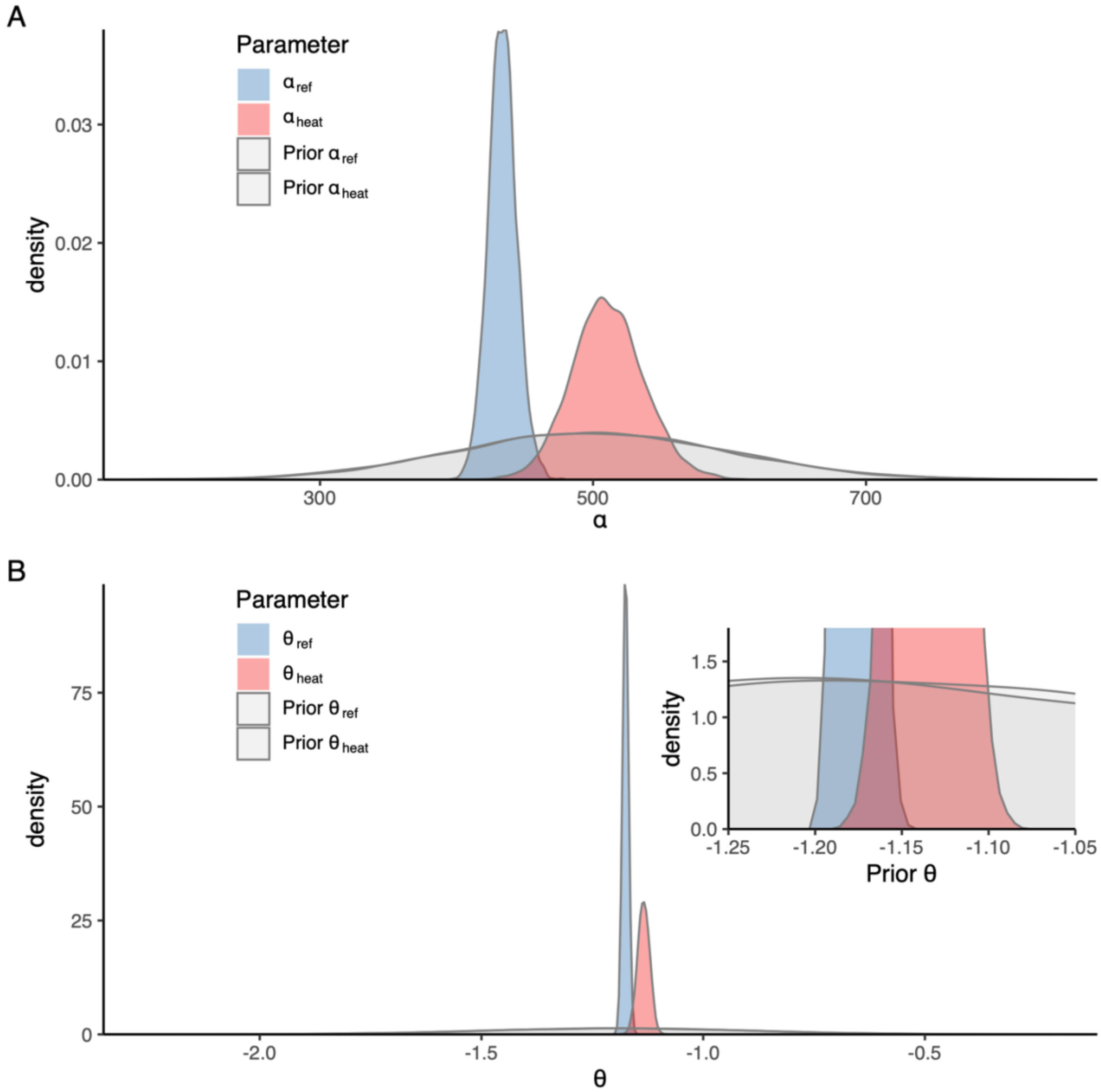


**Fig. S13.** Prior vs posterior distributions for parameters $\alpha$ (A) and $\theta$ (B) in the best allometric growth model (inset in panel (B) is a zoomed-in version to better visualize the priors in the range of the posteriors).

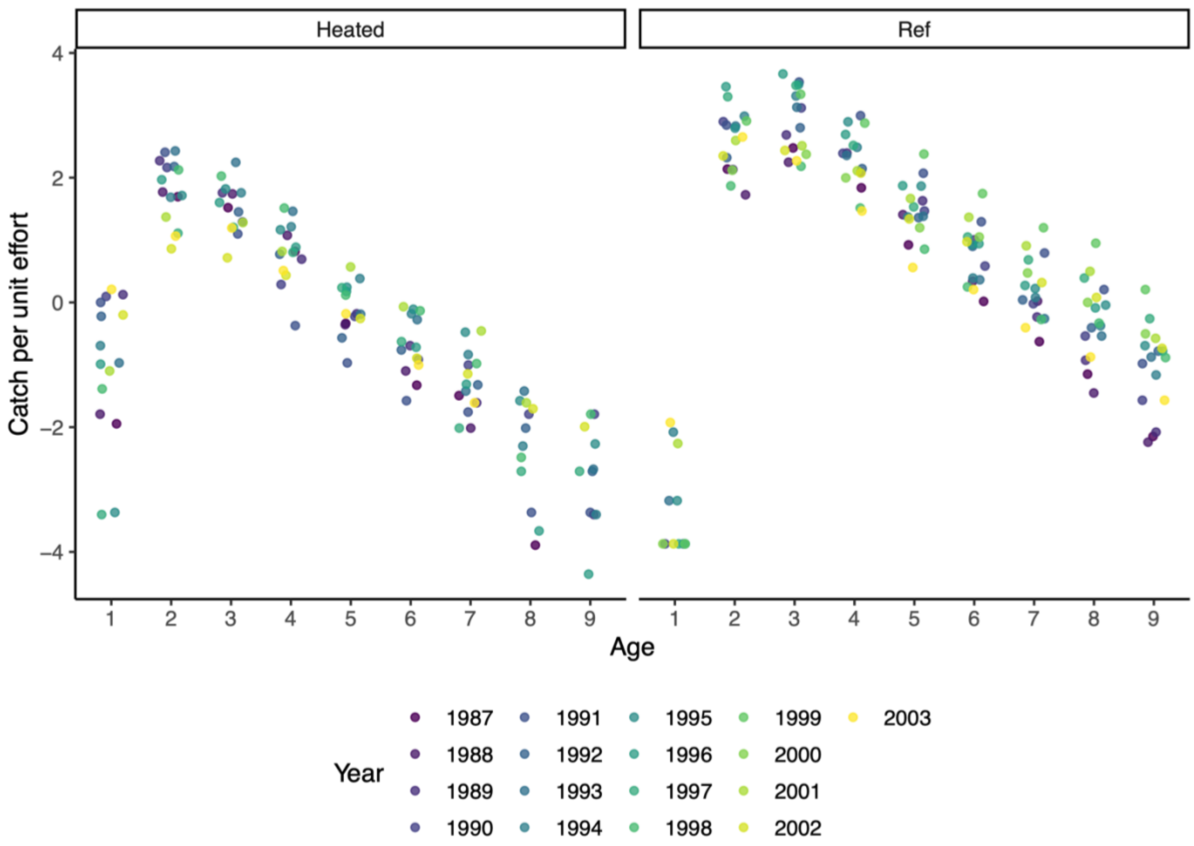


**Fig. S14.** Catch per unit effort (CPUE) as a function of age, by area and year, for determining which ages are representatively caught by the fishing gear. Since the CPUE starts to decline for fish older than 2 years, we selected only fish aged 3 or older for the catch curve regression analysis. Colors indicate catch year.

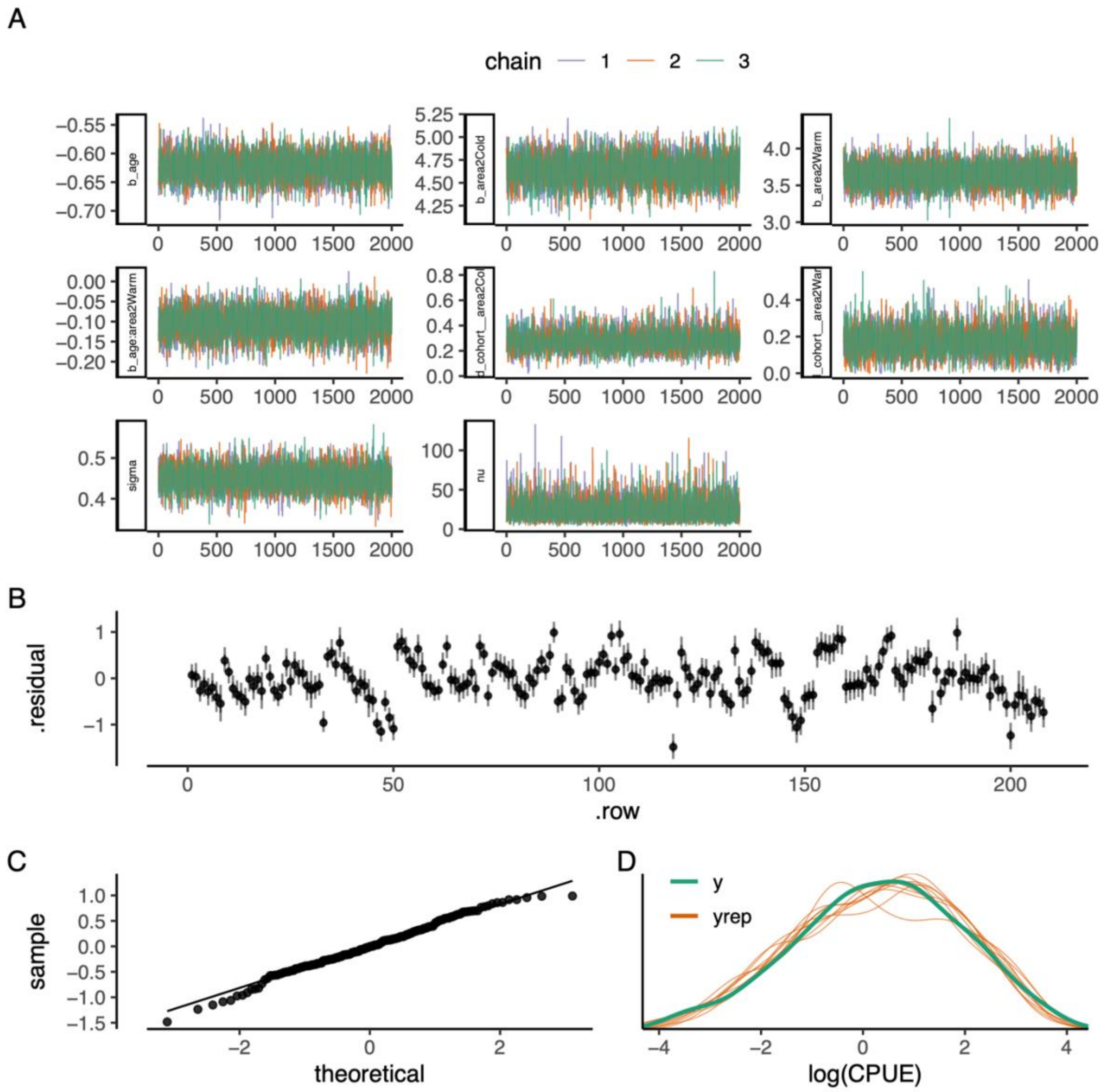


**Fig. S15.** The best catch curve model: (A) traceplot to illustrate chain convergence for key (population-level) parameters, (B) residuals, (C) QQ-plot and (D) posterior predictive check (D).

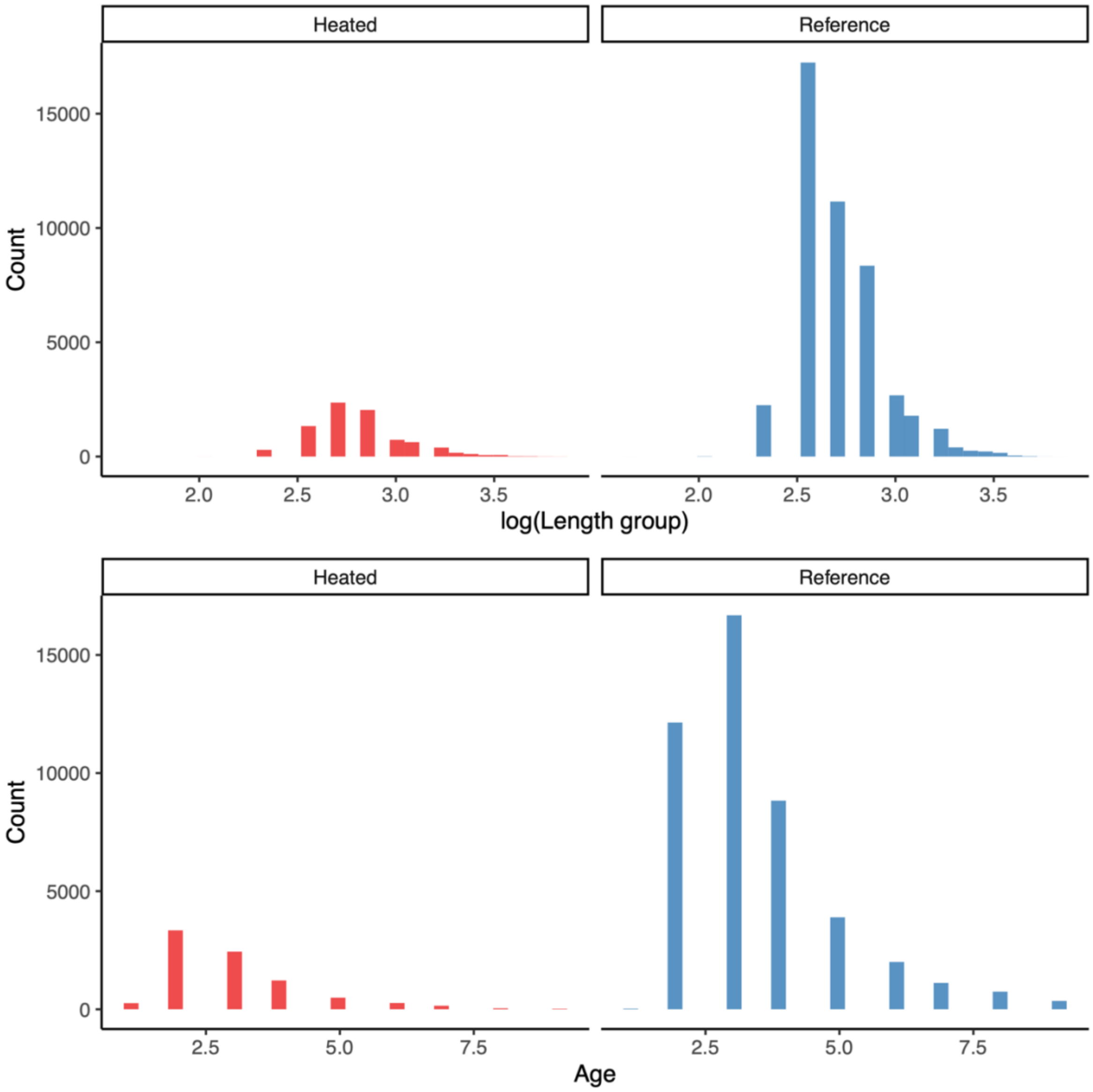


**Fig. S16.** Size (top) and age (bottom) distribution of catches, all years pooled, as used in the lognormal model estimate mean size and catch. Heated area is shown in the left column (in red) and the reference area in the right (blue).

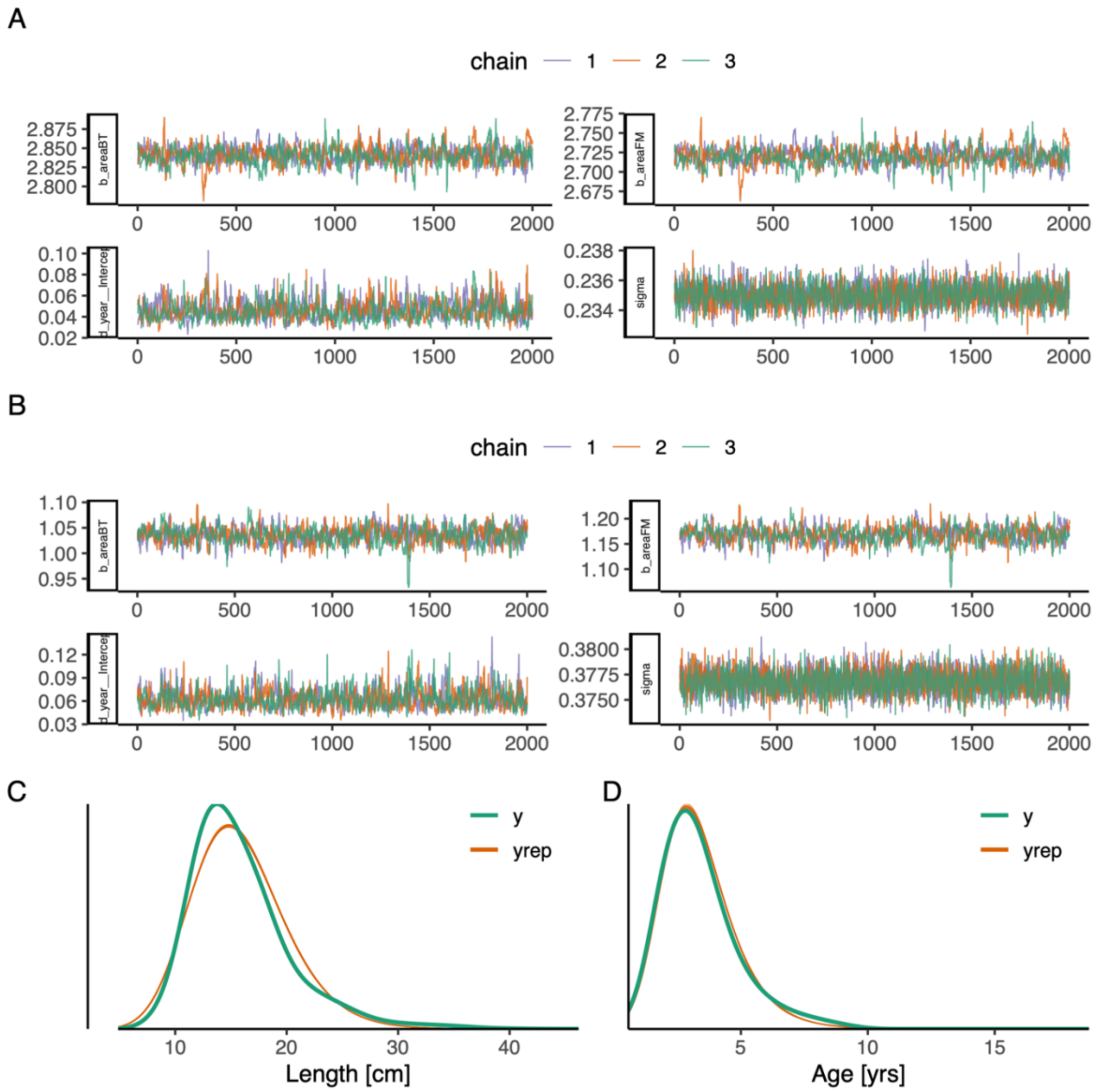


**Fig. S17.** Lognormal length and age models model diagnostics and fit. (A-B) traceplot to illustrate chain convergence for key (population-level) parameters in the lognormal length and age models (respectively), (C-D) posterior predictive checks for the length and the age mode, respectively.
